# *In vivo* nuclear envelope adaptation during cell migration across confining embryonic tissue environments

**DOI:** 10.1101/2025.06.05.658077

**Authors:** Hanna-Maria Häkkinen, Soraya Villaseca, Zain Alhashem, Szymon Chomiczewski, Maxime Desevedavy, Solene Leleux, Atrin Hamidzadeh, Filomena Gallo, Dina El-Zohiry, Victor Petre, Elena Scarpa

**Affiliations:** Department of Physiology, Development and Neuroscience, University of Cambridge, Downing Street, Cambridge, CB2 3DY, United Kingdom; Microscopy Bioscience Platform, University of Cambridge, Downing Street, Cambridge, CB2 3DY, United Kingdom

**Author notes:** these authors contributed equally to the work (Soraya Villaseca, Zain Alhashem). equal contribution (Szymon Chomiczewski, Maxime Desevedavy, Solene Leleux).

## Abstract

In physiology and in disease, cells often migrate through narrow spaces, such as leukocytes undergoing diapedesis or cancer cells during dissemination. Cultured cells under physical confinement have been shown to experience mechanical stress due to deformation of the nucleus. Nuclear deformation can lead to loss of nuclear integrity and DNA damage, and it has been proposed to underlie cancer initiation and progression. *In vivo*, the consequences of physical confinement on physiological, developmental cell migration remain so far unexplored. Here, we use the zebrafish neural crest as an *in vivo* model to address how multipotent embryonic cells respond to physical confinement during developmental migration. By measuring the extracellular space from head to tail along the embryonic body axis, we found that the level of tissue scale confinement increases along the antero-posterior axis. We found that neural crest experience dramatic nuclear deformation during their migrating between adjacent tissues, which quantitatively scales with tissue confinement. By using complementary genetic and mechanical strategies to ablate the surrounding tissue, we observe a rescue of nuclear deformation *in vivo*. Surprisingly, we found that while deformation of the NC nucleus causes stretching of the nuclear envelope, it does not cause DNA damage even upon extreme deformations. Instead, cells adapt, by decreasing both their DNA damage and LaminB2 levels upon entering confined spaces. In summary, we establish the neural crest as a physiological framework uncovering a dynamic adaptation to tissue confinement *in vivo*.

## Introduction

*In vivo*, migrating cells often navigate complex tissue environments, experiencing physical confinement exerted by surrounding cells and tissues and by the extracellular matrix. The ability to negotiate passage through small tissue spaces is essential in physiology. For example, dendritic cells patrol peripheral tissues ^1^ and circulating white blood cells extravasate on demand into tissues upon injury or infection ^2^. In disease, cancer cells often disseminate through confining tissue environments ^3^, and the ability to survive physical confinement correlates with poor prognosis ^4^.

Nuclear deformability represents a limiting factor for cells migrating through small pores ^5^ as the nucleus is considered the stiffest organelle of a cell in adult tissues ^6, 7^. Indeed, when confronted with pores of varying sizes, immune cells can use their nucleus as a gauge to measure their environment to identify the path of least resistance ^8^. The nucleus can act as a proprioceptive device to for cells to sense their surroundings, and can mount functional responses to adapt mode of cell migration, cytoskeleton organisation ^9, 10^ and even chemokine receptor profile ^11^ according to spatial constraints. However, confined cell migration through pores smaller than a critical 3μm threshold ^11^ can lead to compromised nuclear envelope integrity either due to transient ruptures of the nuclear envelope ^12, 13^ which can result in accumulation of DNA damage ^12–16^. This has been shown to drive transition to invasive cancer phenotypes ^16, 17^ and tumour heterogeneity ^14^. In addition, nuclear deformation in absence of ruptures can also induce DNA damage by increasing replication stress ^18^, and accumulation of DNA damage can in turn induce NE ruptures ^19^. Thus, extreme spatial constraints to cell migration can have deleterious outcomes for cells. However, the consequences of confinement on physiological cell migration remain largely unexplored. Recent *in vivo* work in *Drosophila* immune cell migration highlights increased endogenous DNA damage as immune cells move through highly confining haemolymph vessels ^20^. Mouse primordial germ cells, which navigate *in vivo* through diverse tissue environments during their developmental cell migration, also appear to suffer striking cell and nuclear deformations and accumulate significant DNA damage ^21^. However, most immune cells are relatively short-lived and terminally differentiated ^22^ whilst germ cells, being totipotent, are subject to a strict quality control and elimination of unfit cells ^23^.

Whether multipotent cells, essential for embryonic development, can adapt to physical stress in an *in vivo* physiological tissue context has not yet been addressed.

To tackle this question, we used the Zebrafish neural crest, harnessing the natural variation in the neural crest microenvironment to interrogate how migratory cells respond to tissue confinement *in vivo*. Neural crest cells are a migratory, pluripotent stem cell population, forming at the dorsal aspect of the neural tube spanning all the way from head to tail of vertebrate embryos ^24^. Neural crest cells are endowed with invasive and migratory capabilities, and actively migrate long distances across tissues, eventually differentiating into multiple cell types, including cartilage and bone, connective tissue, pigment cells, neurons and glia of the peripheral nervous system. Across vertebrates, cranial neural crest (cNCs) migrate under the ectoderm through large interstitial spaces filled with loosely organised extracellular matrix ^25–27^. In contrast, trunk neural crest (tNC) migrate in dense tissue environments, initially invading narrow inter-tissue spaces between the spinal cord and the somites using a highly conserved migratory pathway ^28–32^. Here, we show that neural crest cells encounter a fine-grained gradient of *in vivo* tissue-scale confinement along the antero-posterior axis of the embryo, which results in dramatic nucleus deformations. Using quantitative *in vivo* imaging, and *in vivo* genetic and mechanical manipulations we find that nucleus shape changes are caused by confinement induced by the surrounding somite tissue. To our surprise, we uncover that neural crest cells do not experience nuclear envelope ruptures. Instead, they adaptively decrease their levels of endogenous DNA damage as well as the levels of LaminB2 when migrating in confined *in vivo* tissue environment.

## Results

### Neural crest nucleus shape behaviours differ across the AP axis of the embryo

To investigate how cells navigate thorough physically changing tissue environments *in vivo*, we used the Zebrafish neural crest migration as a model system. Zebrafish neural crest migrate across a variety of tissue contexts along the antero-posterior axis of the embryo^33, 34^. Cells navigating complex tissue environments deform the nucleus to overcome physical obstacles or to move through dense extracellular matrix^5^, and use this organelle as a gauge to find the path of least resistance ^8^. We asked whether nuclear deformation also occurs in a physiological context during neural crest cell migration through complex *in vivo* tissue environments. We compared cranial neural crest (cNC), anterior trunk neural crest (atNC) of the vagal streams (somites 1-2), trunk neural crest (tNC, somites 7-12) and posterior trunk neural crest (posttNC, somites 27-33, Figure 1A). In zebrafish, cNCs form lateral to the neural keel, from where they disperse into a loose collective of mesenchymal cells ^25, 33^. On the other hand, trunk neural crest cells (tNCs) form a premigratory cohort dorsal to the neural tube from where they start a collective migration as a chain of cells ^35^ between the surrounding trunk tissues (Fig1B). To monitor nuclear shape changes upon cell migration in their native tissue environment, we carried out in vivo live imaging of embryos expressing the *sox10:mG* tissue specific reporter ^35^, which allows visualisation of neural crest cell and nucleus shapes. To visualise the surrounding tissue context, we injected fluorescently labelled dextran ^36^. We observed that, whilst cNC and atNC nuclei maintain a circular, isotropic shape during their migration, midtNC and posttNC dramatically deform their nuclei (Fig 1A’, Movies 1-4). By segmenting nuclear shapes and measuring nucleus circularity and aspect ratio over time we found that each cell experiences multiple deformation events during its migration (Fig S1A,A’). Because nuclear shape changes occur at different timepoints for every cell, to quantitatively compare nuclear morphometrics over time across the cNC, atNC, midtNC and posttNC population we identified the maximum deformation event per each cell track as the timepoint of minimum circularity (Fig S1A, arrow, S1B) and we found that nuclear shape changes become increasingly stronger along the antero-posterior axis of the embryo. (Fig S1B’). Similarly, we found that the duration of each deformation event also increases along the antero-posterior axis of the embryo (Fig S1C). We observed that, whilst premigratory tNC nuclei appeared circular in shape across the tNC population (Fig 1A’, cyan arrowheads, Fig1B) their deformation started as they moved to the migratory zone squeezing under the somite (Fig1A, yellow arrows, dotted line). Using the dextran labelling to align nuclear shape tracks to the timepoint when tNC enter the interstitial space underneath the somite, we observed that midtNC and posttNC nuclei deformed significantly when migrating under the somite across the tNC population (Fig1C,D, arrows). In contrast, atNC nuclei did not significantly change shape as they moved under the somite, and their nuclear shape metrics remained similar to cNCs (Fig1C, S1D). We found that nucleus deformation scales along the anteroposterior axis of the embryo and detected markedly lower nuclear circularities and higher nuclear aspect ratios in mid and posttNC (Fig 1C,D, S1B,B’, S1D). In summary, we show that neural crest cells display striking differences in nuclear deformation dynamics across the embryo.

**Figure 1.**
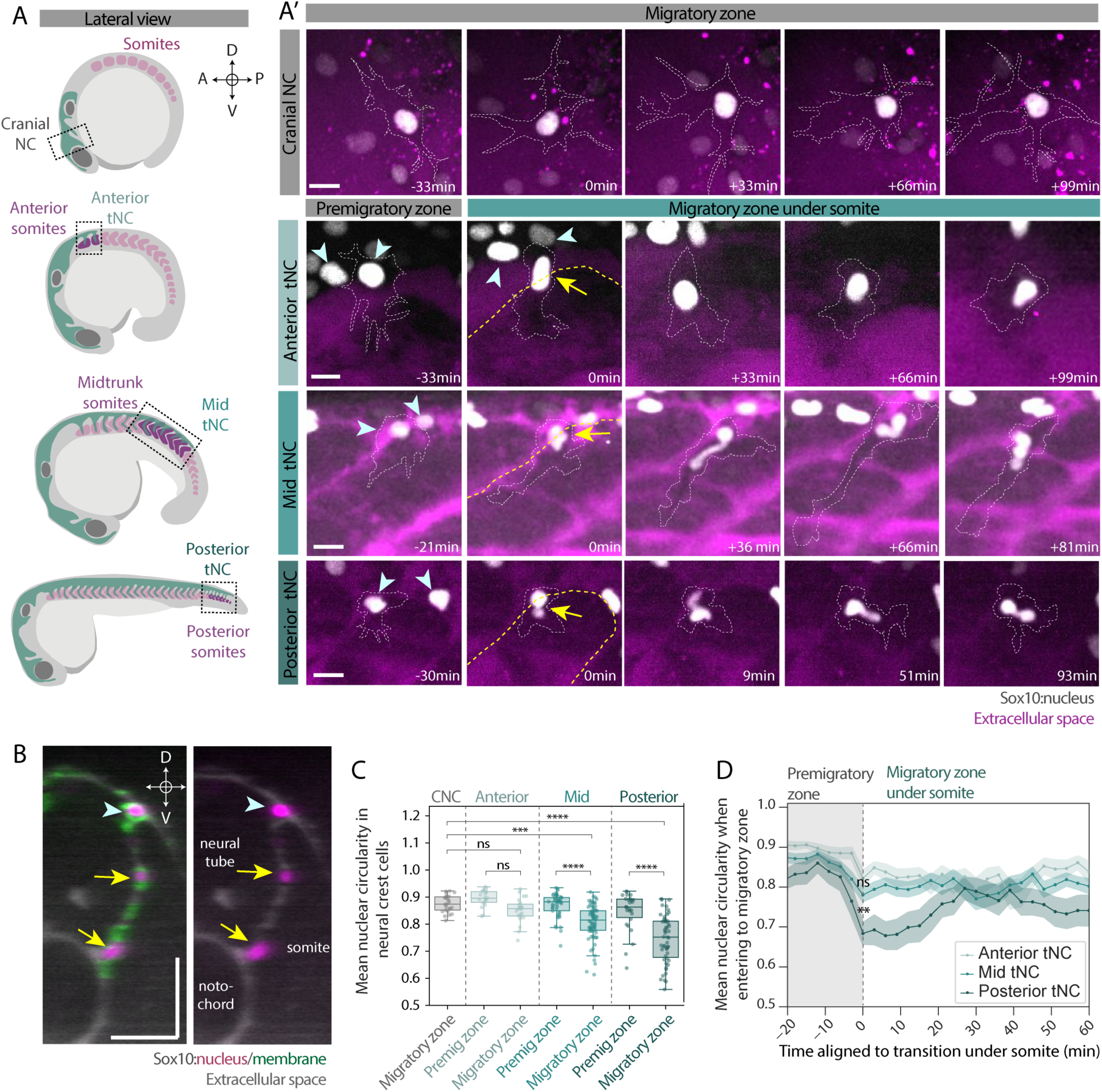
Neural crest nucleus shape behaviours differ across the AP axis of the embryo. A) Sketches of zebrafish embryos at around corresponding developmental stages (cranial 10ss, anterior trunk 14ss, mid trunk 20ss, posterior 30ss) when neural crest (NC) starts migrating from the dorsal region towards the ventral. A’) Representative time series of single neural crest cells during the transition from premigratory zone to the migratory zone in live embryos. Yellow arrowheads indicates the nuclear shape upon the migratory transition. White dashed line indicates the cell membrane. Yellow dashed line indicates the location of the somite boundary in the trunk regions. B) Representative dynamic resliced cross section (line width 10px) at mid trunk region during neural crest migration in live embryo. Light blue arrowheads shows the neural crest still in the premigratory C) Mean nuclear circularity per nucleus in neural crest cells in premigratory region and migratory region along the anterior-posterior body axis. CNC n=25 cells from 5 embryos, Anterior premigratory n=19 cells, anterior migratory n=23 from 9 embryos, mid trunk premigratory n= 45, mid trunk migratory n=72 from 11 embryos, posterior premigratory n= 26, posterior migratory n=51 from 10 embryos. Kruskal-Wallis Dunn’s multiple comparison test *** < 0.005, **** <0.0001. Boxplot shows the median, 1st and 3rd quartile. Each dot represents mean circularity value of a single nucleus over time. D) Mean nuclear circularity aligned to the transitioning from premigratory region to the migratory region under the somite. Anterior n=19 cells from 9 embryos,, Mid trunk n=40 from 11 embryos, Posterior n=23 cells from 9 embryos. Mean and SEM presented. At each time point at least 17 nuclei included. A=anterior, P=posterior, D=dorsal, V=ventral Scale bar 10µm (A’), 40µm (D).

### Tissue-scale confinement differs across the AP axis of the embryo

We reasoned that the different nuclear shape behaviours we observe might be caused by differences in the size and matrix composition of the inter-tissue spaces encountered by the migrating neural crest. Indeed, whilst cranial neural crest migrate lateral to the neural tube into the sparsely organised head mesoderm ^25^ (Fig 2A), trunk neural crest migrate confined in between closely apposed surrounding tissues (Fig 2B), which are lined with a basal membrane of fibronectin and laminin ^37, 38^. To quantitatively measure the size of interstitial spaces, we visualised interstitial spaces ether with injected 10kDa fluorescent dextran or with fluorescently tagged secreted Neurocan-GFP mRNA, which binds to hyaluronic acid ^39^ (Fig S2A). Measurement of interstitial spaces in embryos co-injected with 10kDa dextran and Neurocan-GFP showed near-equivalent values (Fig S2B,C) and thus these approaches were used in parallel. We found that cNCs migrate into a fluid filled cavity posterior to the eye (Fig 2C, C’). By comparing the diameter of the neural crest nucleus as an estimation of its size (Fig S2D) to the width of the inter-tissue space across the mediolateral axis of the embryo (76±23.9 μm (Mean/SD), Fig 2C-D, Fig S2E) we found that the cavity is wide relative to the size of individual cNCs (Fig 2C, arrowheads). Hence, we concluded cNCs migrate in a non-confined tissue environment. Next, we investigated the space along the migratory path in the trunk. *In vivo* imaging of 10kDa Dextran revealed that the spaces along the tNC migratory path, in between the neural tube, somite, and notochord, were extremely narrow(Fig 2B,E-E’). Because we observed that tNC nuclear deformation consistently initiated as tNC nuclei squeezed under the somite (Fig 1C,D), we measured the width of the inter-tissue space at the entry of the neural tube/somite interface (Fig2F, S2F). We found anterior trunk inter-tissue spaces (anterior trunk 4.1±1.2μm (Mean/SD),(Fig 2F)) to be slightly smaller than the mean atNC nuclear diameter (Fig S2D), while mid trunk and posterior trunk inter-tissue spaces were significantly smaller (mid trunk 2.5±0.6μm, 2.8±0.8 μm (Mean/SD)posterior trunk) than midtNC and posttNC nuclear diameter (Fig 2F, S2F) suggesting that midtNC and posttNC nucleus were under constriction during migration. We also found that Neurocan-GFP localized to the inter-tissue spaces in both cranial and trunk regions supporting that inter-tissue space were filled with hyaluronic acid across the AP-body axis (Fig S2B, Fig 2C,C’). As networks of fibrillar collagen can constitute an obstacle for nuclei of migrating cells in 3D ^5^ we next asked whether collagens are also involved in nuclear deformations in neural crest cell migration. Most collagens are not expressed in early zebrafish embryos, however Col1a1a has been previously linked to tNC migration^40^. Immunostaining of Col1a1a (FigS2G) revealed that whilst cranial, anterior and mid-trunk tissues were similarly devoid of type 1 collagen prior to migration onset, posterior trunk tissues displayed detectable levels of Col1a1a around the area where premigratory cells sit in inter-tissue spaces. The presence of Col1a1a in the posterior trunk tissue might explain the stronger nuclear shape changes detected also in the premigratory population in posttNC (Fig 1C,D). In summary, we find that, along the zebrafish trunk, tissue-scale confinement tightly scales with nuclear deformation across migrating neural crest subpopulations.

**Figure 2.**
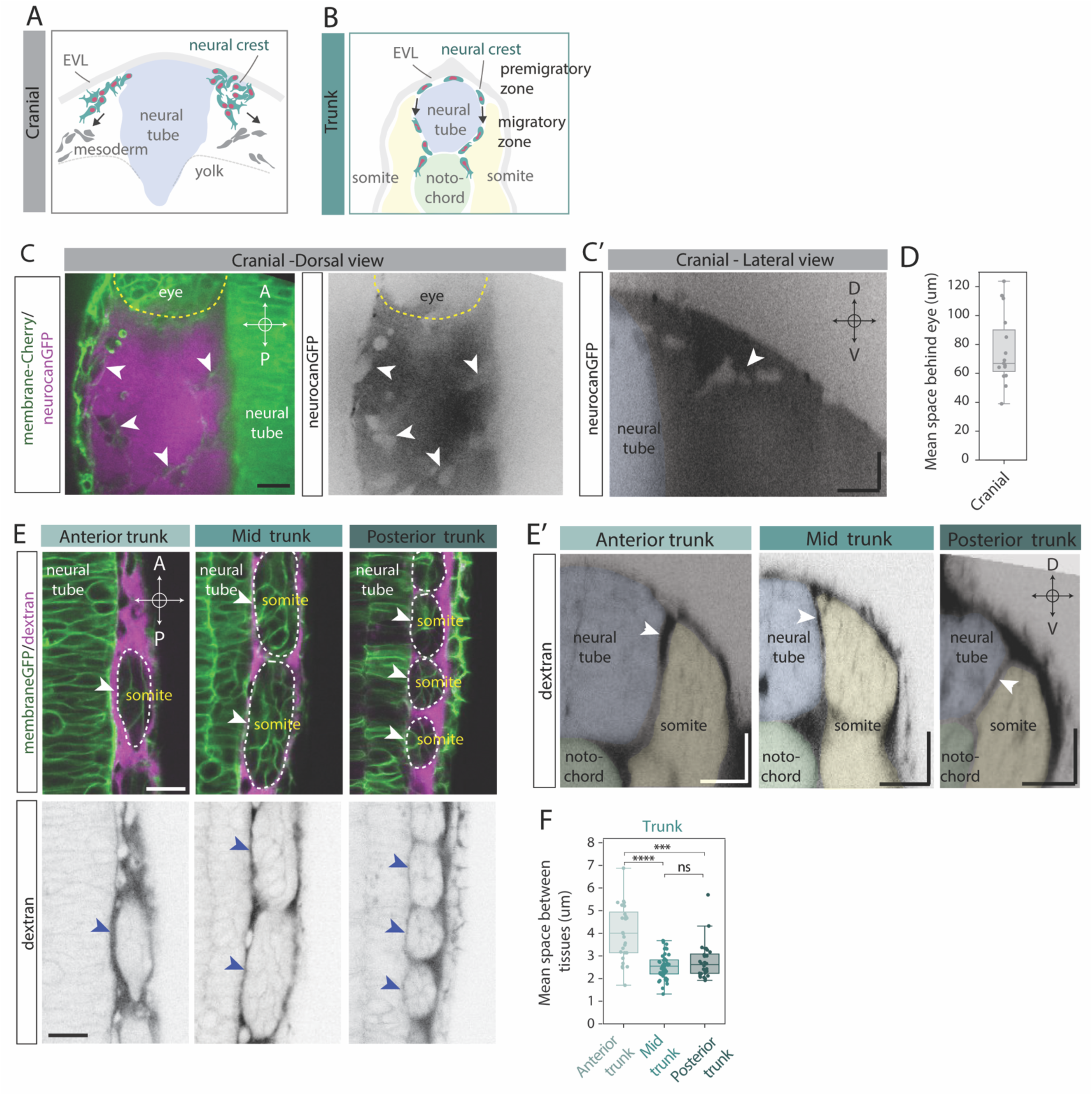
Tissue-scale confinement differs across the AP axis of the embryo. A) Sketch of crossection of cranial region in zebrafish embryo at time of neural crest migration. Arrows indicates the direction of neural crest migration. B) Sketch of crossection of trunk region in zebrafish embryo at time of neural crest migration. Arrows indicates the direction of neural crest migration. C) Representative dorsal view (C) and resliced cross section (C’) of cranial region showing the extracellular space between membrane labelled tissues along the migratory path and the migrating neural crest cells (white arrow heads) in live embryo. Yellow dashed line indicates the location of the eye. D) Mean space between EVL and neural tube behind the eye 10um along the migratory path of CNCs. n=15 measurements, 3 measurements per embryo, 5 embryos.Boxplot shows the median, 1st and 3rd quartile. Each dot represents mean value per embryo. E) Representative dorsal view (E) and resliced cross section (E’) of anterior trunk, mid trunk and posterior trunk at the onset of neural crest migration showing the extracellular space between tissues along the migratory path of neural crest (white (top panel) and blue (bottom panel) arrow heads) in live embryo. White dashed lines shows the location of somites in dorsal view. Pseudo colours indicates the position of adjacent tissues. F) Mean space between a somite and neural tube at different anterior-posterior regions along the first 10um along the neural crest migratory path. Anterior trunk n=28 somites from 8 embryos, Mid trunk n=44 somites from 9 embryos, Posterior trunk n=28 somites from 8 embryos. Kruskal-Wallis Wallis Dunn’s multiple comparison test, ***< 0.0005, ****<0.0001.A= anterior, P=posterior, D=dorsal, V=ventral. Scale bars 20 μm in all panels.

### In vivo perturbations of tissue-scale confinement restore nucleus circularity

ts migrate under the somite through restricted spaces smaller than the resting size of their nucleus, and this correlates with the onset of strong nuclear shape changes. To directly test whether squeezing between the adjacent neural tube and somite tissues is causing the neural crest nuclear deformations, we genetically perturbed tissue-scale confinement in the zebrafish trunk by using the spadetail (*spt*) mutant, where mesoderm tissue is defective and somites do not form in the mid-trunk region ^41^. In *spt* mutants, tNC migrate in an unrestricted manner (Fig 3A) due to reduced slow muscle tissue (Fig S3A)^42^. We reasoned that inter-tissue spaces between the neural tube and the somite tissue remnant in *spt* mutants might be wider than in wild type embryos. Injection of Neurocan- GFP mRNA and quantification of the width of inter-tissue spaces at the neural tube/somite interface indeed revealed the spaces along the tNC migratory path to be significantly wider in *spt* mutant embryos (Fig 3B, green arrows, Fig 3C). We observed that the dorsolateral space between the EVL and the somite was also increased and some tNC could invade this space^43^(purple arrow). To monitor tNC migration, we generated *sox10mG; spt* mutant embryos and labelled inter-tissue spaces using 10kDa fluorescent dextran. We carried out *in vivo* live imaging to compare nuclear shape changes with those of control siblings (Fig 3D). In *spt* mutant embryos, a larger fraction of tNC tend to migrate along the dorsolateral path underneath the EVL ^43^. These cells were excluded from our analysis as they colonised a different inter-tissue niche from control siblings. Instead, we specifically monitored nuclear shape changes for those neural crest cells transitioning from the premigratory area to the inter-tissue spaces between neural tube and somite (Fig 3B, blue arrow, Fig 3D) We found that, in contrast to control sibling embryos, nuclear circularity and aspect ratios was unchanged in *spt* mutants when tNC migrated under the somite remnant (Fig 3E, S3B, Movie 5). These findings suggest that the nucleus deformations of migrating tNC are caused by tissue scale confinement imposed by the surrounding somite tissues. However, as in *spt* mutants mesoderm formation is defective from very early stages of development ^44–46^, we also developed an acute mechanical perturbation of somite formation to independently complement our *spt* findings. We used femtosecond pulsed IR-laser ablation in *sox10:mG* embryos to physically remove somite tissue. This method has been previously validated in Zebrafish and used at similar developmental stages to target the notochord and the tailbud progenitors ^47, 48^. We precisely ablated a portion of presomitic mesoderm at 14-17ss (Fig 3F, S3C), preserving the premigratory neural crest, neural tube and notochord (Fig 3G). This resulted in formation of a gap in the somite segmental organisation, whilst existing somites remained anterior and new somites formed posterior to the ablated region (Fig 3H, S3C). Injection of 10kDa dextran in ablated embryos highlighted a reduction in somite tissue on the ablated side of the embryo, and a marked increase in inter-tissue space, whilst somites on the contralateral side of the embryos were not affected (Fig S3D,E). To monitor nuclear shape changes, we carried out *in vivo* live imaging of *sox10:mG* in ablated embryos starting around 2 hours after ablation when the promigratory to migratory transition took place at the ablated region. Measurement of nuclear shape changes upon transition to the inter-tissue spaces in adjacent, unaffected somites revealed a strong decrease in nucleus circularity (Fig 3H-J, Movie 6)) and a corresponding increase in aspect ratio (Fig S3F). In contrast, cells in the ablated region showed a more subtle decrease in nuclear circularity (Fig 3H-J), and nucleus aspect ratio remained unchanged (Fig S3F). Taken together, we demonstrate, using *in vivo* orthogonal genetic and mechanical perturbations, that tissue-scale confinement originating from the somites imposes nuclear deformations to tNC during developmental cell migration.

**Figure 3.**
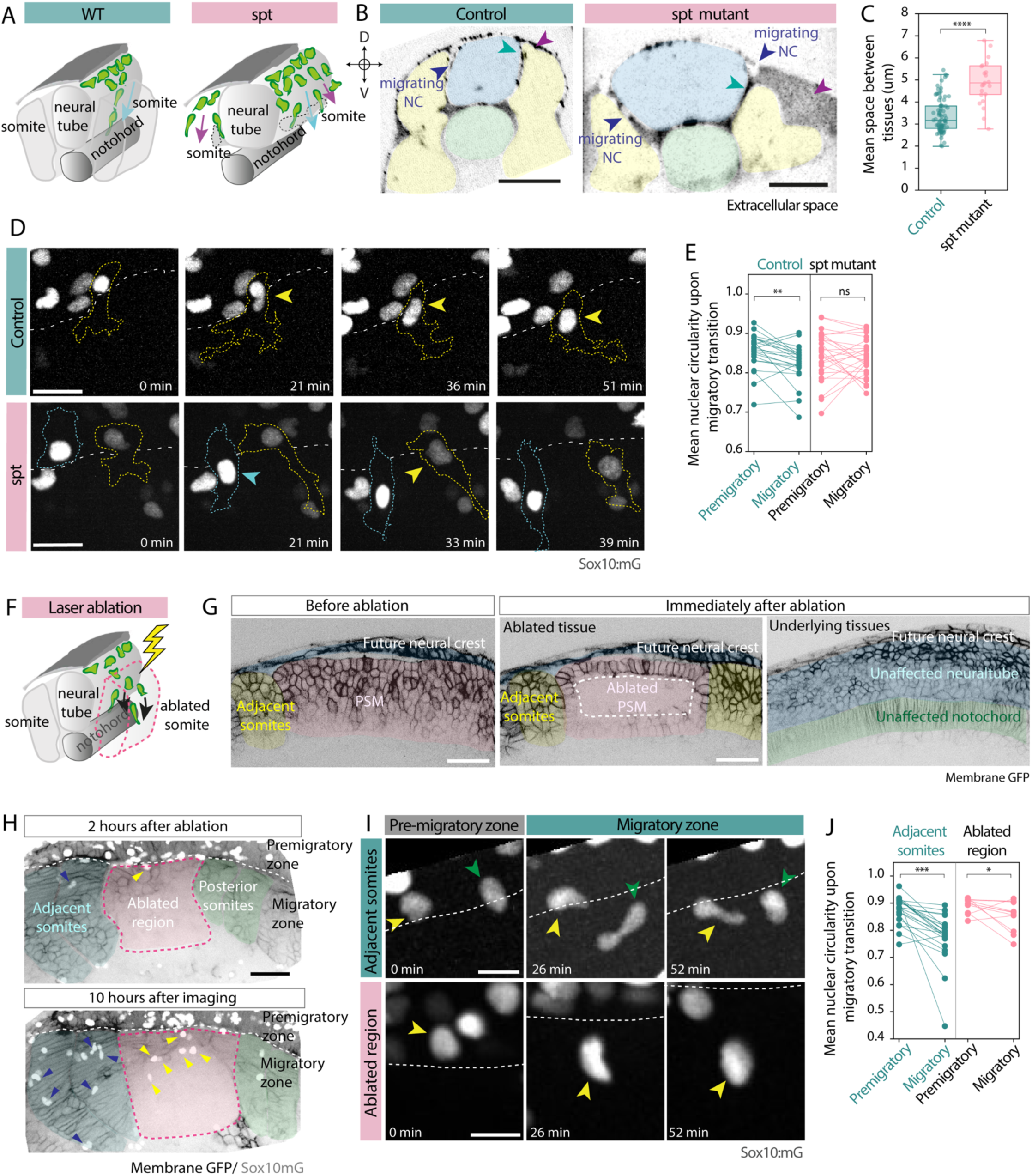
In vivo perturbations of tissue-scale confinement restore nucleus circularity. A) Sketches of tissue organization in zebrafish mid trunk in wildtype (WT) and spadetail mutant (*spt*) fish. Blue arrow heads indicate the neural crest migration through the dorsal path between somite and neural tube, purple arrow head indicates the lateral path between the somite and EVL. Dashed line shows the reduced somite size in mutant line. B) Representative resliced cross section of a mid trunk region in control and *spt* mutant embryo showing the extracellular space between tissues along the neural crest migratory path (green arrow heads) in live embryo. Purple arrow heads highlight the space along the lateral migratory path between EVL and somite. Pseudo colours facilitates the visualization of adjacent tissues. C) Mean width of extracellular space along the dorsal path in control and *spt* mutant embryos. Control n=80 somites from 9 embryos, spt mutant n=23 somites from 8 embryos. Mann-Whitney test, ****<0.0001. Boxplot shows the median, 1st and 3rd quartile. Each dot represents a mean width of the extracellular space along dorsal path per each somite. D) Representative time series of neural crest cells during the transition from premigratory zone to the migratory zone in control and in spt mutant embryo. Arrowheads indicates the nuclear shape upon the migratory transition. Yellow and blue dashed line shows the location of cell membranes during migration. White dashed line shows the location of the somite boundary in the trunk regions. E) Mean nuclear circularity before and after transitioning to the migratory zone. Control n=22 cells from 2 embryos, spt mutant n=26 cells from 4 embryos, Wilcoxon matched-pairs signed rank test **<0.005. F) Sketch of laser ablation approach to remove individual somites. Black arrows shows the direction of the neural crest migration. Dashed line shows the ablated somite. G) Representative images of membraneGFP labelled presomitic mesoderm (PSM) and intact surrounding tissues before and immediately after laser ablation. Pseudo colours indicates the location of surrounding tissues. Dashed line highlights the ablated region. H) Representative images of tissue organization and neural crest cell location at the onset of migration (upper panel) and 10hours after imaging (lower panel) in laser ablated embryo 5 hours after the ablation. Global membrane staining visualise the location of intact somites. Blue arrow heads highlights the neural crest migrated in intact adjacent region while yellow arrow heads highlight neural crest migrated into the ablated region. Dashed line separates the premigratory zone from the migratory zone. Pseudo colours indicates the location of the intact and ablated tissues. Maximum projection of neural crest channel overlayed with single plane of membraneGFP channel for better visualization. I) Representative time series of single neural crest cells during the transition from premigartory zone to the migratory zone in intact adjacent regions and ablated regions of the embryo. Arrowheads indicates the nuclear shape upon the migratory transition. White dashed line separates the premigratory region form the migratory region. J) Mean nuclear circularity before and after transitioning to the migratory zone at intact adjacent regions and ablated regions of the embryo. Adjacent region n=21 cells from 4 embryos, ablated region n=12 cells from 4 embryos, Wilcoxon matched-pairs signed rank test *<0.05, ***<0.0001.Scale bars 40 μm (B, G), 20 μm (D, H), 10 μm (I).

### Confined migration of tNC results in nucleo-cytoplasmic leakage

Our results show that during developmental migration in narrow inter-tissue spaces, tNC undergo striking, repeated nuclear deformations. Cells are prone to nucleus envelope (NE) ruptures *in vitro* when forced through pores smaller than 3μm ^3, 11–16^. To ask whether the dramatic nucleus deformations observed in migrating tNC lead to NE ruptures, we focused on tNC at mid trunk region, as we observed the maximum difference between tissue space and nucleus size in this region (Fig 2E-F, S2D). To interrogate NE ruptures, we live imaged tNC cells expressing a nuclear localised emerald-GFP (nls-emGFP) under control a neural crest specific Gal4 ^49^. To validate that nls-emGFP could be used as a proxy for nuclear envelope integrity in zebrafish, we imaged tNC cell divisions (Fig S4A). We observed a strong increase of nls-emGFP fluorescence in the cytoplasm upon NE breakdown at mitotic entry (Fig S4A, arrow), followed by sequestration of nls-emGFP within the nucleus upon mitotic exit (Fig S4A, yellow arrowheads). Using nls-emGFP as a reporter for NE integrity, we carried out in vivo live imaging of migrating tNC (Fig 4A, Movie 7) and premigratory tNCs (Fig 4B, Movie 8). Quantification of nls-emGFP nuclear and cytoplasmic intensity throughout deformation revealed leakage of small fraction of nls-emGFP into the cytoplasm and simultaneous decrease of nuclear nls-emGFP intensity upon maximum nuclear deformation (Fig 4C,C’). nls-emGFP relocalization was not observed in premigratory tNC, where nuclear deformation events were subtle (Fig 4D,D’). In premigratory tNC cells nuclear nls-emGFP intensity slightly increased over time, possibly due to an increase in activity of the sox10 promoter over developmental time ^50^, whilst the cytoplasmic nls-emGFP fraction remained constant (Fig 4D’).

**Figure 4.**
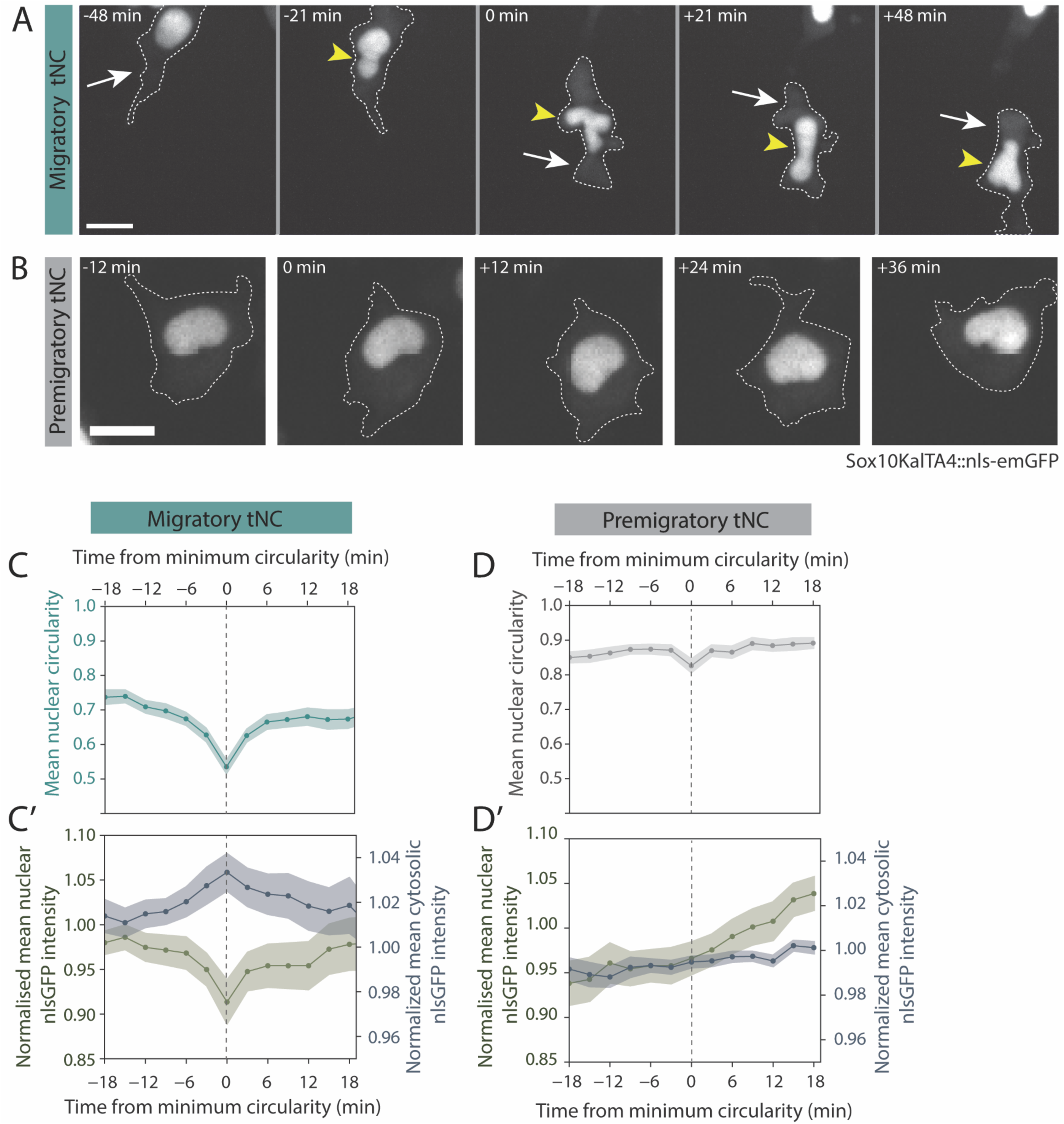
Confined migration of tNC results in nucleo-cytoplasmic leakage. A) Representative time series of nuclear deformation and nls-emGFP localization in migrating neural crest cell in mid trunk region. Yellow arrowheads highlights the nuclear shape during migration. White arrows indicates the cytosolic nls-emGFP before, during and after the maximum deformation of the nucleus. Dashed line indicates the cell membranes. B) Representative time series of nuclear deformation and nls-emGFP localization in migrating neural crest cell in cranial region. Dashed line indicates the cell membranes. C) Mean nuclear circularity during minimum circularity event during trunk neural crest migration (C) and normalised nls-emGFP intensity in nucleus and cytosol aligned to the minimum circularity event (B’). n=50 cells from 9 embryos, Mean and SEM presented. At each time point at least 29 nuclei included. D) Mean nuclear circularity during minimum circularity event during cranial neural crest migration (C and normalised nls-emGFP intensity in nucleus and cytosol aligned to the minimum circularity event (D’). n=15 cells from 4 embryos, Mean and SEM presented. At each time point at least 10 nuclei included. Scalebar 10 μm in all panels.

To further substantiate these observations, we monitored nls-emGFP nucleus-cytoplasmic leakage in cNCs, which migrate in an unconfined manner in a large interstitial cavity (Fig 2A,2C-D). We found that, like premigratory tNC, cNCs did not show a clear decrease in nls-emGFP nucleus intensity nor an increase in the cytoplasmic intensity upon nucleus shape changes (Fig S4B-C’). These findings suggest that the confined migratory TNC cells, but not their premigratory counterparts or cNCs, their non-confined counterparts, show a transient loss of NE integrity. However, quantification of nls-emGFP only showed a 5% decrease in nuclear intensity, accompanied by a correspondingly small increase in cytoplasmic intensity in migrating tNC. Upon NE ruptures, chromatin becomes exposed to the cytoplasm and can be detected by cytoplasmic DNA binding proteins, such as BAF^13, 51^. For this reason, we further investigated nuclear envelope ruptures using BAF-mCherry^13, 51^. As previously reported in human cells ^52^, we were able to visualise a transient, bright accumulation of BAF at it cross-bridges mitotic chromosomes into a single nucleus in late anaphase (Fig S4D). However, we were not able to identify BAF positive foci in proximity of the nucleus even during most dramatic nucleus deformation events (Fig S4E, Movie 9), suggesting that the nucleocytoplasmic leakage of nls-emGFP could be a result of nuclear pore stretching ^53, 54^, rather than of a loss of NE integrity. To test this possibility, we carried out scanning electron microscopy (SEM) of wild type Zebrafish trunk cross-sections to visualise the nuclear envelope of confined tNCs in detail. SEM revealed an intact nuclear envelope in migrating tNC (Fig S4F-F’’, arrows). Measurement of nuclear pore size revealed that a significant increase in migrating tNCs compared to premigratory tNCs (Fig S4G). Together, our results support that during trunk neural crest migration, dramatic nuclear deformation events lead to stretching of the NE pores, causing a transient nls-emGFP leakage without compromising the NE integrity.

### *In vivo* confined migration of tNC does not lead to accumulation of DNA-double strand breaks

Our results show that during developmental migration, neural crest cells dramatically deform the nucleus, which results in passive leakage of the nls-emGFP across the nucleo-cytoplasmic barrier, likely caused by stretching of the nuclear pores rather than by NE rupture. In cultured cells, it is well established that nuclear envelope ruptures can result in accumulation of DNA damage. This has been reported to be caused by several mechanisms. DNA damage can follow a long lasting nucleus envelope rupture event, which can cause the cytoplasmic nuclease TREX1^55^ to enter the nucleus ^16^or the sequestration of DNA repair factors away from the chromatin due to their leakage into the cytoplasm ^14, 15, 56^. In addition, nuclear deformation, in absence of nuclear envelope ruptures, can cause DNA damage by inducing stalling of DNA replication forks in cells in the S-Phase of the cell cycle ^18^. Trunk neural crest initiate migration during S-phase ^57, 58^. Because we observe striking nucleus deformations in tNC upon initiation of migration (Fig 1D), we asked whether these deformation events could result in DNA damage. We immunostained *sox10:mG* embryos for phosphorylated γH2AX to visualise DNA double strand breaks ^59^. Surprisingly, we observed that tNC migrating under the somite have significantly lower levels of γH2AX fluorescence intensity compared to premigratory tNC (Fig 5A, compare pink arrows and arrowheads, Fig 5B). We hypothesised two possible scenarios: either migrating tNCs are able to rapidly repair DNA double strand breaks, or tNC do not accumulate *de novo* DNA double strand breaks when confined. To address this, we measured the dynamics of DNA damage foci formation and resolution in live embryos. We generated mRNA encoding a truncated mApple-53BP1, which localises to DNA double strand breaks and is optimised for *in vivo* live imaging ^60^. To validate this reporter in the zebrafish, blastula stage embryos were treated with the PARP inhibitor Olaparib ^59, 60^ to induce DNA double strand breaks, which were readily detected as 53BP1 subnuclear foci (Fig S5A, arrows). To ask whether migrating tNC experience DNA damage as a consequence of *in vivo* confined cell migration, we mosaically injected mApple- 53BP1mRNA in sox10KalTA4::nls-emGFP embryos. This allowed us to concomitantly monitor nuclear shape changes, 53BP1 foci lifetime and fluorescence intensity fluctuations. We found that 53.8% of tNC did not show any distinct 53BP1 foci during migration. For the fraction of cells that did form one or more detectable foci during live imaging, we observed that these appeared irrespective of nucleus shape prior to foci formation for both premigratory and migratory tNC (Fig 5C,D, Movie 10, Movie 11). Also, foci lifetime was in the timescale of minutes for both premigratory and migratory tNCs (Fig S5B). In mammalian cells, it 53BP1 intensity can increase with a delay of up to 2 hours relative to the nucleus squeezing event ^12–16^. To directly interrogate whether DNA damage could occur upon in vivo confinement, we measured 53BP1 intensity specifically in cells that transitioned from the premigratory area to the small inter-tissue space under the somite, and measured 53BP1 foci number for 2 hours before and after the under somite transition (Fig 5E). Whilst we could detect a sharp change in nucleus shape upon transition (Fig 5E, arrow), we found that 53BP1 foci number was very low throughout nuclear deformation events when tNC migrated under the somite (Fig 5E). Since most cells did not form any detectable 53BP1 foci, we also measured the ratio between the maximum and mean fluorescence intensity to detect transient, more dynamic, fluctuations in 53BP1 signal. Similarly to foci quantifications, we observed that 53BP1 intensity remained constant (Fig 5F) despite dramatic nucleus deformation upon under somite, and even slightly reduced after the under somite transition (Fig 5F’). Consistent with our live observations, we confirmed no correlation between tNC nucleus circularity and endogenous γH2AX intensity (Fig S5C). Overall, our results show that in migrating, confined tNCs, endogenous DNA damage levels are lower than in premigratory tNC. Using *in vivo* live imaging, we find that the lifetime of DNA damage foci in live cells is similarly short in both premigratory and migratory tNC, suggesting that the lower levels of γH2AX in migrating tNC are not due to faster DNA repair. We observed that transition of tNCs into the highly confined under somite space does not lead to foci formation or to an increase in 53BP1 intensity peaks despite strong nucleus shape changes. We conclude that tNC do not accumulate DNA damage during *in vivo* confined cell migration.

**Figure 5.**
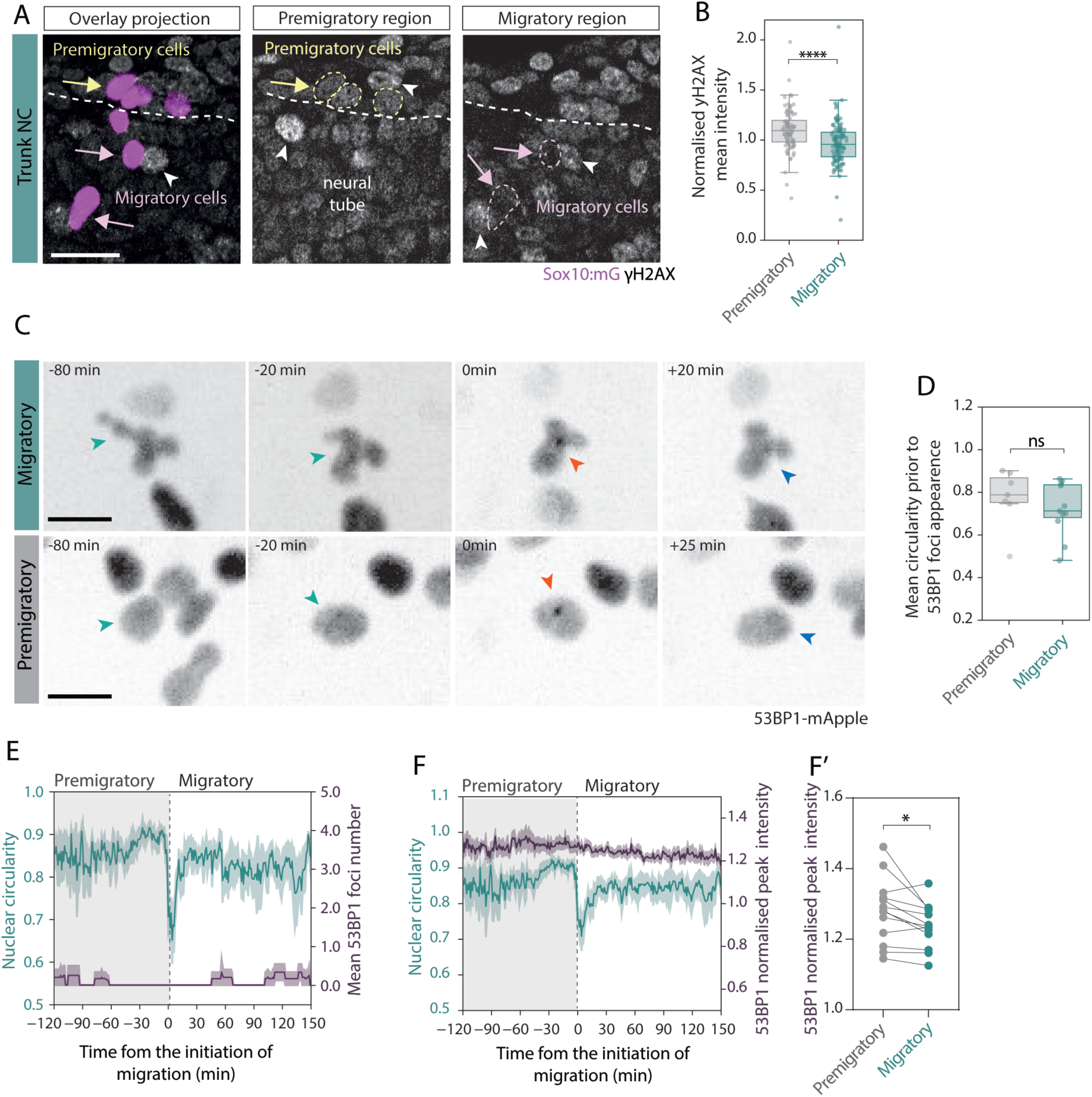
In vivo confined migration of mid TNC does not lead to accumulation of DNA-double strand breaks. A) Representative maximum projections and single z-slices showing of yH2AX localization in mid trunk region in premigratory (yellow arrows) and migratory neural crest cells (purple arrows) in fixed embryo. Dashed lines indicates nucleus boundaries, white arrow heads highlights the intensity levels in surrounding tissues. B) yH2AX intensity in premigratory and migratory cells normalised to surrounding tissues. Premigratory cells n=89 cells from 7 embryos, migratory cells n=131 cells from 7 embryos. Kruskal-Wallis test, ****<0.0001. Boxplot shows the median, 1st and 3rd quartile. Each dot represents a normalised mean value of a nucleus. C) Representative time series of 53BP1 localization in migratory and premigratory neural crest cells in mid trunk region. Cyan arrowheads shows the nuclear shape before the 53BP1 foci appearance, red arrowhead indicates the 53BP1 foci in the nucleus, and blue arrowheads indicates the nuclear shape after the foci disappears. D) Mean circularity up to 60min before the appearance of 53BP1 foci in premigratory and migratory cells. Premigratory n=7 foci from 5 cells and migratory n=11 foci from 8 cells, from 5 embryos. Mann-Whitney test, ns>0.05. Boxplot shows the median, 1st and 3rd quartile. Each dot represents a foci. E) Nuclear circularity over time aligned to the transition from premigratory region to migratory region together with mean number of 53BP1 foci in foci positive cells. n=8 cells from 6 embryos. Mean and SEM shown. F) Nuclear circularity over time aligned to the transition from premigratory region to migratory region together with normalised 53BP1 peak intensity and (F’) normalised 53BP1 peak intensity per nucleus before and after premigratory to migratory transitioning. n= 13 cells from 8 embryos. Paired t-test, *<0.05. Scale bar 20 μm (A), 10 μm (C).

### The neural crest nucleus dynamically adapts to confinement

Our results show that tNC do not accumulate DNA damage as a result of confined cell migration, instead, endogenous DNA damage levels appear reduced in tNCs migrating under tissue confinement. This suggests that the migrating trunk neural crest nucleus may dynamically adapt to confinement to reduce stress. To test this hypothesis, we investigated the lamin composition of neural crest nuclei by immunostaining for LaminA/C, LaminB1 and LaminB2. As previously reported ^61^ early zebrafish embryos do not express lamin A/C except for a small subset of somite cells (Fig S6A, top inset). LaminB1 is expressed at low levels in the CNS and at very low levels in neural crest cells (Fig S6A, bottom inset), whilst it is more highly expressed in somite tissue (Fig S6A). However, we found LaminB2 to be ubiquitously expressed in the early zebrafish embryo, including in neural crest cells. Quantification of LaminB2 levels at the nuclear periphery reveals that tNCs significantly reduce LaminB2 upon under somite migration (Fig 6A,B), thus suggesting that the nuclear lamina composition adaptively responds to confinement. To resolve dynamic changes of the nuclear lamina and its associated inner nuclear envelope partners during *in vivo* confined migration, we mosaically injected mRNA encoding the LEM-domain protein Lap2β-EGFP ^10^ in sox10-Kalt4 embryos, where chromatin is labelled with H2B-mCherry ^62^ (Figure 6C, Movie 12). High spatiotemporal resolution of migrating tNC cells revealed that Lap2β-EGFP is dynamically depleted from the front of the nucleus, resulting in a polarised distribution of Lap2β-EGFP at the nuclear envelope during nuclear deformation events, which recovers after deformation (Fig 6D, Movie 12). In contrast, in non-confined migrating cNCs, Lap2β-EGFP does not show a polarized localisation (Fig S6B, C). Lap2β links chromatin to the nuclear periphery ^63, 64^ and mediates force transmission from the nuclear lamina to chromatin upon stretching^65^. Hence, reduced levels of LaminB2, together with localised reduction of Lap2β may reduce tethering of chromatin at the nuclear periphery, facilitating the passage of the nucleus trough small tissue spaces and limiting *de novo* accumulation of DNA damage.

**Figure 6.**
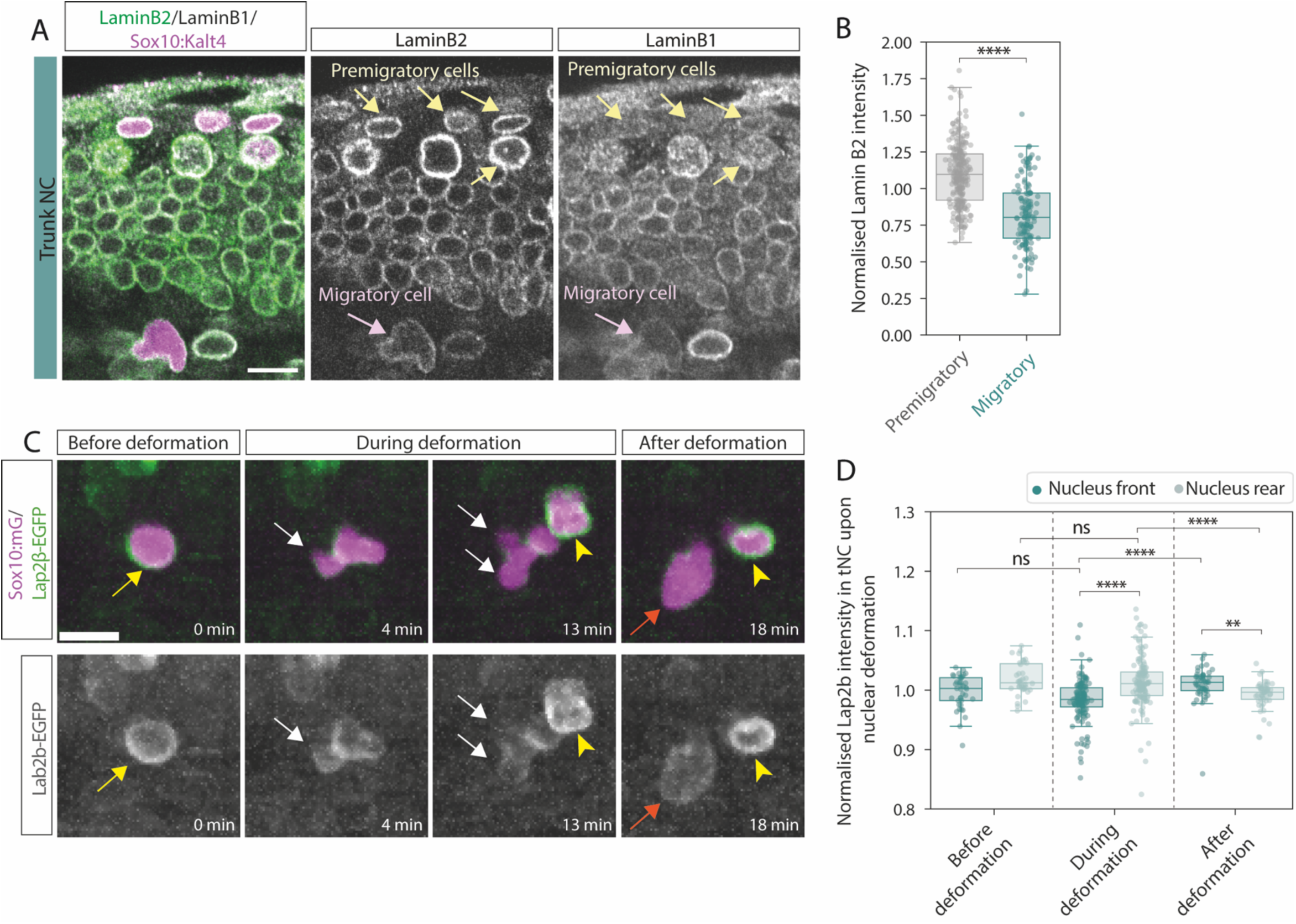
The neural crest nucleus dynamically adapts to confinement. A) Representative images of expression of nuclear laminB2 and B1 in premigratory (yellow arrows) and migratory (pink arrows) neural crest cells in mid trunk region in fixed sample. (max projection of 2 1um Z-stacks) B) Nuclear laminB2 intensity in premigratory and migratory neural crest cells in mid trunk normalised to the mean LaminB2 intensity across tNCs (Premigratory+ Migratory). Premigratory n=244 cells, Migratory n=135 cells from 9 embryos. Kruskal-Wallis Dunn’s multiple comparisons test, **** < 0.0001. Boxplot shows mean, 1st and 3rd quartile. Each dot represents a nucleus. C) Representative images of Lap2β localization in the nucleus of migratory cell in mid trunk region during nuclear deformation event. Yellow arrows shows the Lap2β localization at the leading front of the nucleus before deformation. White arrow heads shows the depletion of lap2b at the bulking side of the deforming nucleus. Orange arrows shows the relocalization of Lap2β at the leading front of the nucleus after deformation. Yellow arrow heads shows Lap2β localisation in neighbouring cell nucleus without deformation. D) Normalised Lap2β intensity at the leading front and the rear of migratory trunk neural crest cell nucleus before, during and after nuclear deformation event. Before deformation n=39 timepoints from 15 cells, during deformation n=132 timepoints from 15 cells, after deformation n=48 timepoints from 15 cells.n= 4 embryos. Boxplot shows mean, 1st and 3rd quartile. Each dot represents a timepoint. Scale bar 10 μm in all panels.

## Discussion

We establish the Zebrafish neural crest as a novel *in vivo* model to investigate the cellular responses of a multipotent stem cell population to tissue scale confinement in a physiological *in vivo* context. Our findings reveal that, during their physiological, developmental migration, neural crest cells invade highly confining inter-tissue spaces across the vertebrate body axis. This results in dynamic, repeated nuclear deformation events, characterised by striking nucleus shape changes that are initiated as tNCs transition from the dorsal premigratory area to the narrow inter-tissue space under the somite. We find that tissue-scale confinement and col1a1a density increase with developmental time along the anteroposterior body axis, and this quantitatively scales with the extent of nuclear deformation experienced by migrating neural crest cells. Strikingly, we observe that in contrast to tNC, the unconfined cNCs and tNC of the vagal streams, which encounter a lesser degree of confinement, do not show significant nuclear deformation. Using a nuclear localised emGFP, we observe nucleocytoplasmic leakage in migrating tNC, but not premigratory tNC or unconfined migrating cNCs. However, our live imaging of BAF-mCherry and SEM measurements of nuclear pore width suggests that nucleocytoplasmic leakage of nls-emGFP might be caused by stretching of the nuclear pores, rather than by NE ruptures (Fig S4G). It is well established that mechanical stress can deform the nucleus ^53^, leading to an expansion in nuclear pore diameter ^66^, which can result in passive leakage of proteins of low molecular weight across the nuclear pore ^53, 54^. Recent work in migrating Drosophila immune cells also found that only a small fraction of cells show NE ruptures after a significant nuclear deformation in an *in vivo* context (Karling and Weavers, 2025) . *In vitro,* NE ruptures consistently occur when immune cells are confined under a PDMS confiner below a critical height of 3μm ^11^. However, these devices actively impose a drastic pressure drop of 10kPa on cells ^67^, whilst reported anisotropic tissue-scale pressure measurements *in vivo* in the zebrafish presomitic mesoderm are around the 0.15-0.25 kPa mark ^68, 69^. While neither the elastic modulus nor tissue-scale pressure measurements in the segmented Zebrafish mid-trunk have yet been reported, these tissues are likely to be stiffer than the mesenchymal, fluid tailbud ^70^.

One possible interpretation of our observations could be that tNC moving physiologically through soft embryonic tissues may not experience sufficient pressure on the nucleus to suffer NE ruptures. However, we find that endogenous LaminB2 strongly decreases upon tNC transition to the under somite spaces. This suggests that tNC cells may actively soften as they encounter a confining environment. In line with this hypothesis, AFM measurements of cytoplasm stiffness in LaminB2 knock-out fibroblasts reveals marked softening and an increased capability for confined cell migration ^71^. Moreover, *Xenopus* neural crest become softer as they transition from a premigratory to migratory state ^72^, hence, fine tuning of LaminB2 levels could allow the nucleus to match the rheological properties of surrounding tissues to facilitate passing of narrow constrictions, thus minimising mechanical stress on the nucleus. Further evidence of an active, dynamic, adaptation of tNC nuclei to migration under confinement is provided by our live quantitative observations of the LEM domain protein Lap2β-EGFP. Lap2β mediates force transmission from the nuclear envelope to chromatin upon mechanically induced cell stretching ^65^. Thus, depletion of Lap2β from the leading edge of deforming nuclei (Fig 6C) might release tensional stresses on chromatin, hence facilitating its passage through small inter-tissue pores ^73^.

Neural crest are long lived stem cells that, upon completion of their migration, give rise to a vast plethora of cell types in the embryo. Neural crest cells colonise many different tissue niches within the body, and their evolution is thought to be instrumental to the success and diversification of vertebrates ^74–78^. Thus, it is likely that this cell type might be endowed with robust mechanisms to withstand stressors encountered during their developmental journey. Indeed, in striking contrast with other instances of *in vitro*^12–14, 16,18^ and *in vivo* confined cell migration^20, 21, 79^ we find that neural crest cells do not accumulate DNA damage when transitioning into a confined environment. Instead, we report that levels of DNA damage, both endogenous (Fig 5A-B) and measured using a live reporter for DNA double-strand breaks (Fig 5F), are reduced in tNC upon *in vivo* confined migration. How do neural crest cells protect their nucleus from mechanical stresses? Possible molecular mechanisms could include BMP and Wnt signalling pathways. These pathways are active throughout the lifetime of neural crest cells ^80^, and govern initiation of tNC migration under the somite by controlling the G1/S cell cyle transition ^81^. Recent work has demonstrated that both BMP and Wnt signalling protect from DNA damage in the regenerating zebrafish heart ^82^ and in human pluripotent stem cells ^83^ by alleviating replication stress, so it is tempting to speculate these signalling pathways may also play a protective role in the pluripotent, S-Phase migrating, tNC. Other possible genome protection mechanisms in migrating tNC might be linked to regulation of cell cycle checkpoints upon initiation of confined migration. Indeed, zebrafish tNC cell cycle progression is regulated by Notch signalling, which prolongs S-Phase and delays G2/M transition of migrating leader cells, so that these predominantly divide mid-way through migration at the notochord/somite interface ^57^. Interestingly, in cultured cancer cells, migration through confinement delays cell cycle progression ^84, 85^.

Why would tNC be endowed with such protection mechanisms? Failure to mount a protective response to migration through confining environment might have deleterious consequences for tNCs, as it might lead to accumulation of mutations and tumour initiation. Importantly, human cancers such as melanoma and neuroblastoma are tNC derived^86, 87^. Polymorphisms predisposing to neuroblastoma occur in genes involved in the control of DNA repair and cell cycle checkpoints pathways ^88^. Neuroblastomas are highly heterogeneous and showing a high degree of copy number variations and chromotripsis, which correlate with poor prognosis and the origin of which is not yet understood ^89^. Growing clinical evidence shows that neuroblastomas originate in utero, at early stages of tNC development ^90^, strongly suggesting that chromosomal instability occurs early in neuroblastoma tumorigenesis and that human tNCs may suffer inherent genomic instability^88, 91^. Clinical studies^92^ also showed that depending on anatomical location human neuroblastomas carry different kinds of chromosomal aberrations. Of note, P53 Zebrafish mutants spontaneously develop tNC derived malignant peripheral nerve sheath tumours^93, 94^. Thus, under unfavourable genetic circumstances the physical environment of tNCs may influence neuroblastoma pathogenesis. On the other hand, as melanomas often hijack embryonic neural crest cell states^87, 95^, re-activation of embryonic adaptability programs could contribute to aggressiveness. Importantly, nuclear adaptation to migration in confined environments by uncoupling of inner nuclear membrane and lamina via Lap1c was recently reported in human metastatic melanoma^4^. In summary, our work provides insight in how an evolutionarily successful, multipotent cell type, the neural crest, adapt to challenging extracellular environments in a physiological tissue context by modulating their nuclear envelope composition, limiting accumulation of DNA damage and thus ensuring robust developmental migration in the early embryo.

## Supporting information

Supplementary movie 1. cNC migration

Supplementary movie 2. Anterior tNC migration

Supplementary movie 3. mid tNC migration

Supplementary movie 4. post tNC migration

Supplementary Movie 5.Trunk neural crest cells migration in spt mutant embryos

Supplementary Movie6. Neural crest migration in laser ablated embryo

Supplementary Movie7. nls-emGFP dynamics in trunk migratory neural crest

Supplementary Movie8. nls-emGFP dynamics in trunk premigratory neural crest

Supplementary Movie9. BAF-mCherry localisation during trunk neural crest migration

Supplementary Movie10. 53BP1-dynamics during trunk neural crest premigratory to migratory transition

Supplementary Movie11. mApple-53BP1-dynamics in premigratory trunk neural crest

Supplementary Movie12. Lap2beta-EGFP dynamics in migratory trunk neural crest cells

## Acknowledgments

HMH is supported by a University of Cambridge Herchel Smith Postdoctoral Fellowship. ES is supported by a Royal Society Dorothy Hodgkin Fellowship (DHF\R1\201118). DEZ was supported by a Royal Society Enhancement Award to ES (RF\ERE\210154). SV and AH are supported by an MRC New Investigator Grant to ES (MR/W024519/1). SL, SC and MD were supported by SmartsUp studentships. VP was supported by a Gatsby Foundation summer studentship. We thank Dr Karin Müller and the Electron Microscopy Facility of the Microscopy Bioscience Platform for technical support and insightful advice. We thank Dr Claudia Linker for generously sharing the *sox10:mG* and *sox10-Kalt4* zebrafish lines and her expertise, time, reagents and knowledge. The Tol2Kit ^96, 97^ was a gift from Prof Koichi Kawakami. We thank Dr Harold Burgess for sharing the 14xUAS-BGI-nucEmGFP-POUT plasmid ^98^, Dr Verena Ruprecht for the gift of the pCS2+-Lap2β-EGFP plasmid, Dr Adam Shellard for the gift of pCS2+-membraneGFP and membrane-mCherry. pCs2+-ss-NCan-GFP was a gift from Dr Kelly Smith. sox10-Lyn-tdTomato plasmid DNA was a gift from Dr Stephan Heermann. BAF-mCherry was a gift from Dr Stephen Royle. *Spt1-b104r* mutant fish were a gift from Dr Ben Steventon. *Sox10:KaltA4* zebrafish were a gift from Dr Peter Arthur-Farraj and Prof David Lyons. We are very grateful to Dr Clare Buckley for initially hosting ES Zebrafish on her PPL (PF2F15847) and we thank Dr Ben Steventon for critical reading of the manuscript.

## Author contributions

HMH, SV, ZA, SC, SL, MD, AH, DEZ, FG and ES carried out experiments. VP tested several segmentation pipelines and contributed a Fiji macro (see Methods) for the annotation of Stardist segmented nuclei. HMH prepared figures, HMH and ES wrote the paper, and all authors commented. ES secured funding.

---

**Supplementary Figure 1.**
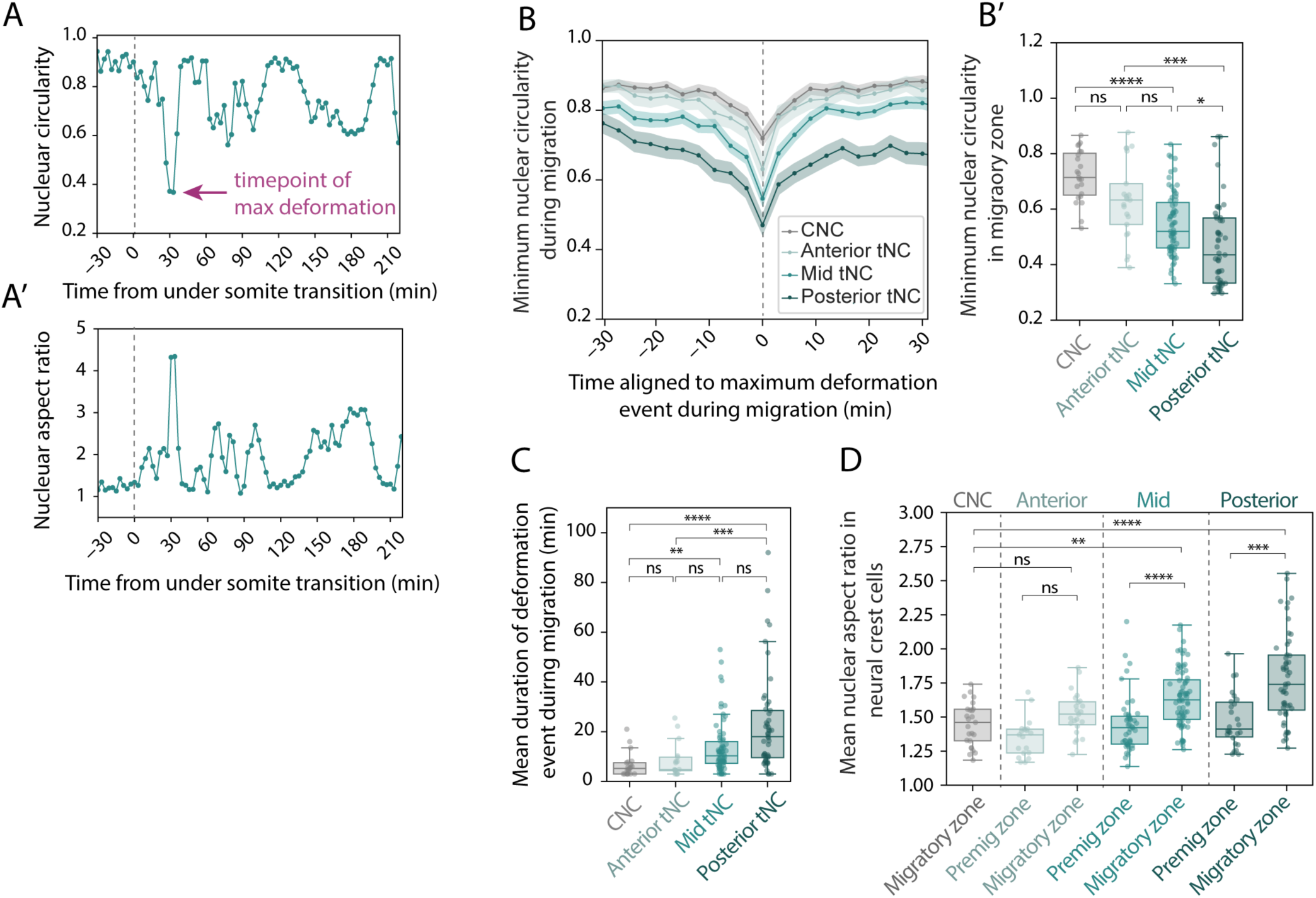
A) Example of nuclear shape fluctuation in a single mid trunk neural crest cell aligned to the transitioning from premigratory region to the migratory region under the somite presented in nuclear circularity (A) and nuclear aspect ratio (A’). B) Mean nuclear circularity aligned to the minimum circularity event during NC migration at different anterior-posterior regions. Mean and SEM presented. At each time point at least 16 nuclei were included, and B’) comparison of the minimum nuclear circularity values in migratory cells. CNC n=23 cells from 5 embryos, N=3, Anterior n = 21 cells from 9 embryos, Midtrunk n=67 from 11 embryos, Posterior n=49 cells from 10 embryos . Kruskal-Wallis Dunn’s multiple comparison test *<0.05, ***< 0.0005, ****<0.0001. Boxplot shows the median, 1st and 3rd quartile. Each dot represents the minimum circularity value of a single nucleus. C) Mean duration of nuclear deformation events per nucleus calculated as consecutive timepoints nucleus circularity remains under a mean circularity value. CNC n=21 cells from 5 embryos, Anterior n=17 cells from 9 embryos, Midtrunk n=70 from 11 embryos, N=3, Posterior n=49 cells from 10 embryos. Kruskal-Wallis Dunn’s multiple comparison test **< 0.005, ***=0.0005, ****<0.0001. Boxplot shows the median, 1st and 3rd quartile. Each dot represents a mean duration of deformations per nucleus with at least one nuclear deformation event. D) Mean nuclear aspect ratio per nucleus in neural crest cells in premigratory region and migratory region along the anterior-posterior body axis. CNC n=25 cells from 5 embryos, Anterior premigratory n=19 cells, anterior migratory n=23 from 9 embryos, midtrunk premigratory n= 45, mid trunk migratory n=72 from 11 embryos, posterior premigratory n= 26, posterior migratory n=51 from 10 embryos. Kruskal-Wallis Dunn’s multiple comparison test test **<0.01, ***<0.0005, ****<0.0001. Boxplot shows the median, 1st and 3rd quartile. Each dot represents mean aspect ratio value of a single nucleus over time.

**Supplementary Figure 2.**
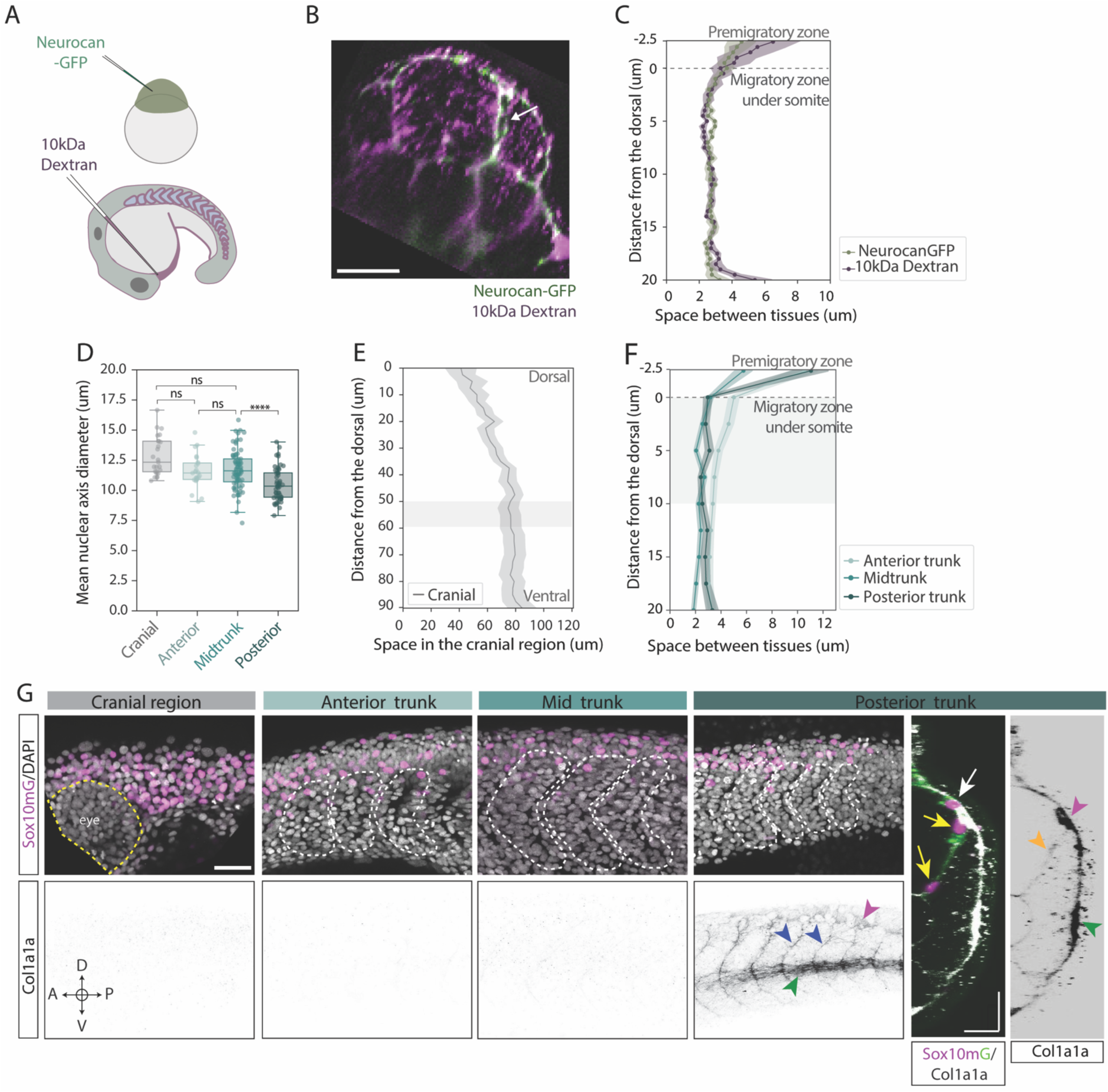
A) Sketch of mRNA injection of Neurocan-GFP at 1-cell stage and Dextran injection prior to imaging. B) Representative dynamic cross section showing the extracellular space between adjacent tissues using dextran and Neurocan-GFP in live embryo. C) Mean width of extracellular space between somite and neural tube using alternative labelling methods for extracellular space, Neurocan-GFP and Dextran, in mid trunk region. n=4 somites from 2 embryos. Mean and SEM presented. D) Mean nuclear axis diameter in neural crest cell at different anterior-posterior region. CNC n=25 cells from 5 embryos, Anterior n=23 cells from 9 embryos, Mid trunk n=77 cells from 11 embryos, Posterior n=53 cells from 10 embryos. Boxplot shows the median, 1st and 3rd quartile. Each dot represents a mean nuclear axis diameter per nucleus over time. Kruskal-Wallis Dunn’s multiple comparison test ****< 0.0001. E) Mean width of extracellular space along the migratory path of neural crest in cranial region from dorsal to ventral behind the eye. Shaded area indicates the region included to the panel Fig2D. n=15 measurements, 3 measurements per embryo, 5 embryos. Mean and SEM presented. E) Mean width of extracellular space in along the migratory path of neural crest in anterior, mid and posterior trunk regions, aligned to the initiation of migratory path. Shaded area indicates the region included to panel Fig2F. Anterior n=28 somites from 8 embryos, N=3, Mid trunk n=44 somites from 9 embryos, Posterior n=28 somites from 8 embryos. Mean and SEM presented. G) Representative images of collagen1a1a in fixed embryos at cranial, anterior trunk, mid trunk and posterior trunk regions. Maximum projections (between 45 to 95 slices with 0.5um z-step) shows the neural crest and surrounding tissues (upper panel) and localization of col1a1a at these regions (lower panel). Blue arrow heads highlights the accumulation of collagen 1a1a at the somite boundaries, green arrow heads highlights the collagen 1a1a at the lateral line. Cross section of the posterior trunk shows the localization of neural crest cells and collagen 1a1a at the premigratory region (white arrow and purple arrow head) and in the migratory path (yellow arrows and orange arrow head). Yellow dashed line indicates the location of an eye in cranial regions. White dashed lines indicates the somite boundaries. A= anterior, P=posterior, D=dorsal, V=ventral. Scale bars 40 μm (B, G; lateral view), 20 μm (G; cross section).

**Supplementary Figure 3.**
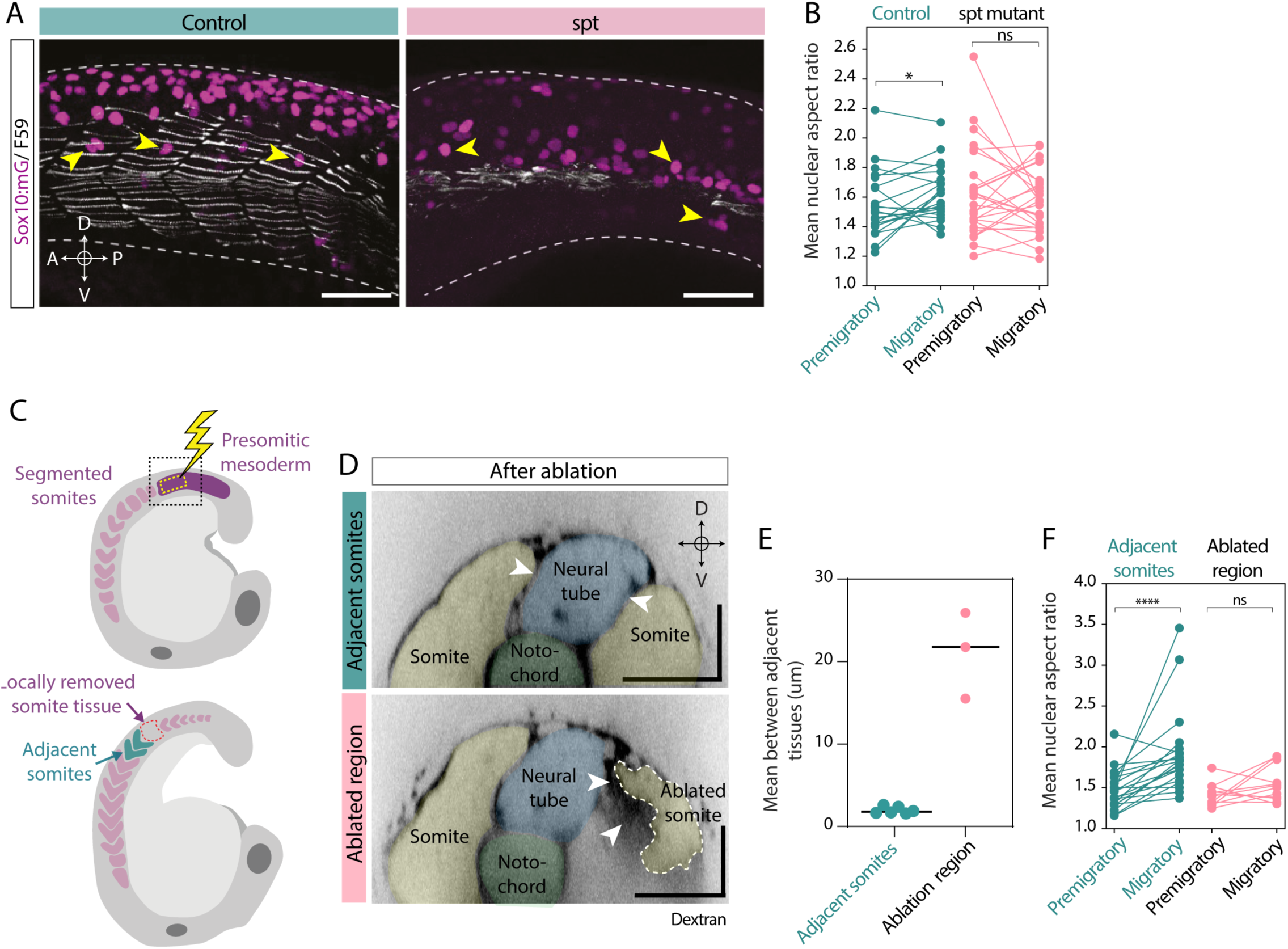
A) Representative images of fixed control and spt mutant embryos at mid trunk region. Maximum projections (n=30 slicez, 1 μm z-step) shows the migrating neural crest cells (yellow arrow heads) and localization of myosin heavy chain (F59) in somite tissue. B) Mean nuclear aspect ratio before and after transitioning to the migratory zone. Control n=22 cells from 2 embryos, spt mutant n=26 cells from 4 embryos, Wilcoxon matched-pairs signed rank test, *< 0.05. Each dot represents a mean value of single nucleus in given region. D) Sketch showing the targeted region witing the embryo during and after laser ablation. D) Representative images from cross sections showing the extracellular space in laser ablated embryos in ablated region and intact somite region. Dashed lines highlights the ablated somites. Pseudo colours indicates the location of adjacent tissues. White arrow heads points the neural crest migratory region. E) Mean space between adjacent tissues along 30um from the initiation of the dorsal migratory path at intact somite regions and at ablated somite region. Adjacent somites n=6 from 3 embryos, ablated region n=3 from 3 embryos. N=1. Mann-Whitney test, * < 0.05. F) Mean nuclear aspect ratio before and after transitioning to the migratory zone at intact adjacent region and ablated regions of the embryo. Adjacent control region n=21 cells from 4 embryos, N=2, ablated region n=12 cells from 4 embryos, N=2, Wilcoxon matched-pairs signed rank test, ****<0.0001. Scale bar 50 μm (A), 40 μm (D).

**Supplementary Figure 4.**
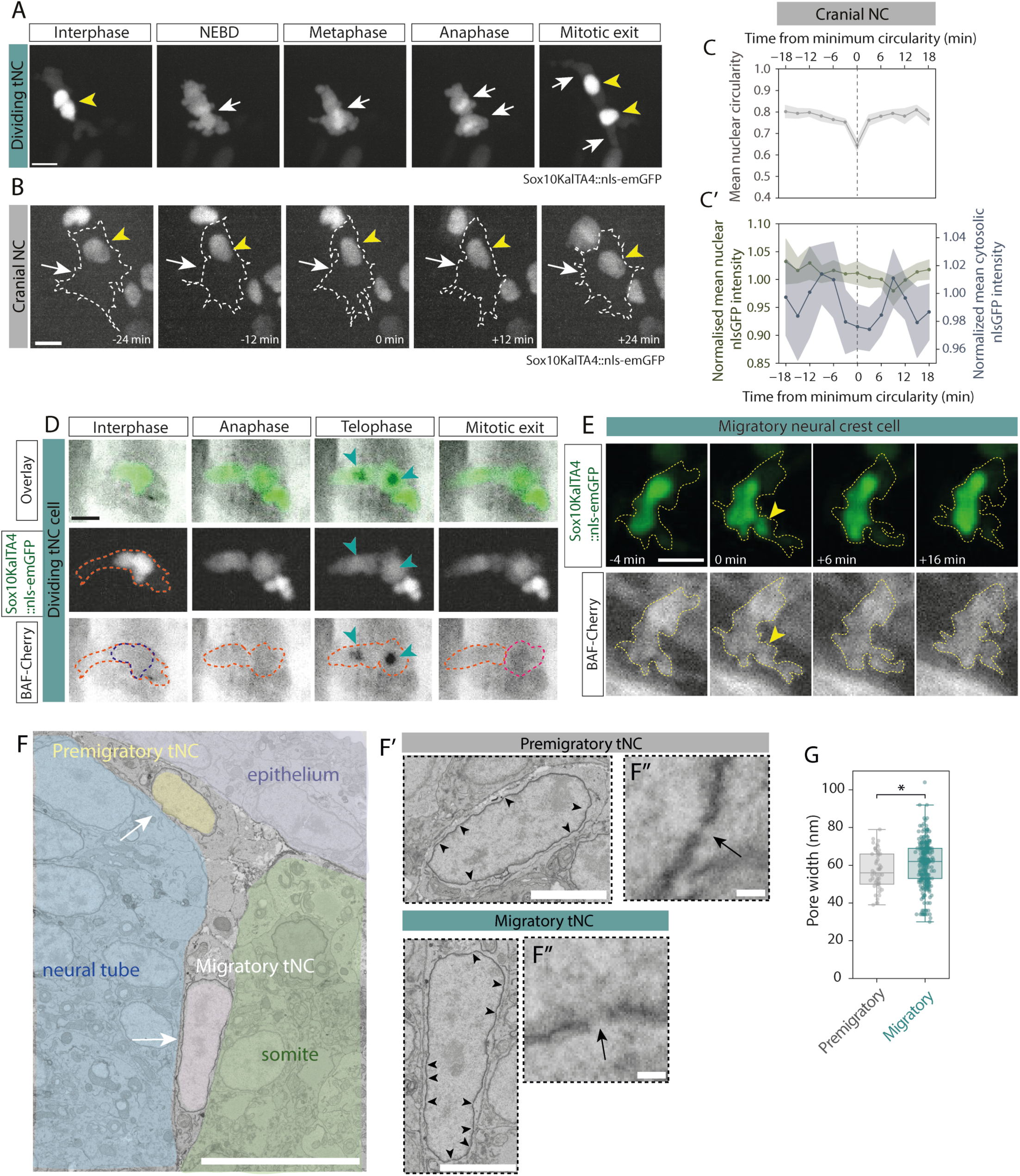
A) Representative timeseries of nls-emGFP localization during cell division in migrating neural crest cell in mid trunk region. White arrow heads shows the depletion of of nls-emGFP in the nucleus during cell division. Yellow arrowheads shows the localization of nls-emGFP in nucleus in interface before cell division and the relocalization in the nuclei in daughter cells after division. B) Representative timeseries of nls-emGFP localization premigratory neural crest cell in mid trunk region. Dashed lines shows the cell membrane localization. Yellow arrowheads highlights the nuclear shape during migration. White arrows indicates the cytosolic nls-emGFP before, during and after the maximum deformation of the nucleus. Dashed line indicates the cell membranes. C) Mean nuclear circularity during minimum circularity event during cranial neural crest migration (C) and normalised nls-emGFP intensity in nucleus and cytosol aligned to the minimum circularity event (C’). n=12 cells from 5 embryos. Mean and SEM presented. D) Representative timeseries of BAF localization during cell division in nls-emGFP positive migrating neural crest cell in mid trunk region. Cyan arrowheads shows the rapid accumulation of BAF in telophase. E) Representative timeseries of BAF localization during nuclear deformation event in nls-emGFP positive neural crest cell during migration in mid trunk region. Yellow arrow heads shows the nuclear shape during maximum deformation event. Dashed outlines highlight the cell shape in both the nls-emGFP and BAF-mCherry. F) Representative image of electron microscopy slice in mid trunk region (F) and representative zoom-in of single migratory and premigratory neural crest cells (F’) and further zoom in for a single NE pore (F’’). White arrows shows the premigratory and migratory cells. Black arrows and arrowheads shows the individual NE pores in the zoom-in images. Pseudo colours facilitates the visualisation of adjacent tissues and neural crest cells, G) NE pore width in premigratory cells and migratory cells. Premigratory n=49, Migratory n=287 pores from 23 sections, all sections from 1 embryo. Mann-Whitney test, * <0.05. Boxplot shows the median, 1s and 3rd quartile. Each dot represents a single pore. Scale bar 10 μm (A, B, D, E, F), 3 μm (F’), 100 nm (F’’).

**Supplementary Figure 5.**
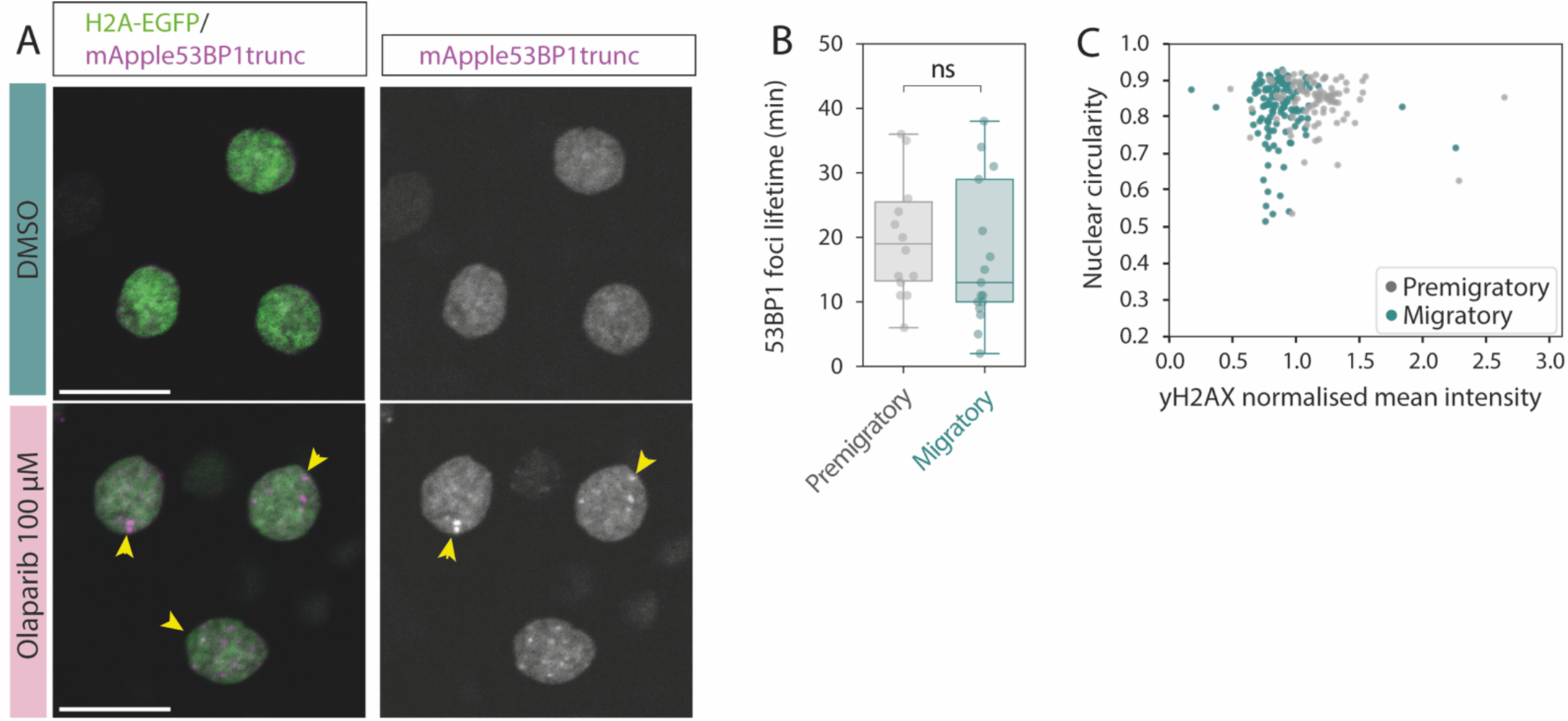
A) Representative images of 53BP1trunk accumulation in nucleus after Olaparib treatment. Yellow arrowheads highlights the localization of 53BP1 foci.n=4 DMSO embryos n=4 Olaparib treated embryos. B) 53BP1 foci lifetime in neural crest cells in mid trunk region during premigratory and migratory phase. Foci in premigratory phase n=14 from 11 cells, foci in migratory phase n=17 foci from 14 cells. Mann-Whitney test, ns>0.05. Boxplot shows the median, 1st and 3rd quartile. Each dot represents a foci. C) Normalised yH2AX intensity over nuclear circularity values in premigratory and migratory trunk neural crest cells. Premigratory n=89 cell, 7 embryos.Spearman R=0.08 p=0.39 (ns), migratory n=131 cells, 7 embryos. Spearman R=-0.03 p=0.77 (ns). Scale bar 20 μm.

**Supplementary Figure 6.**
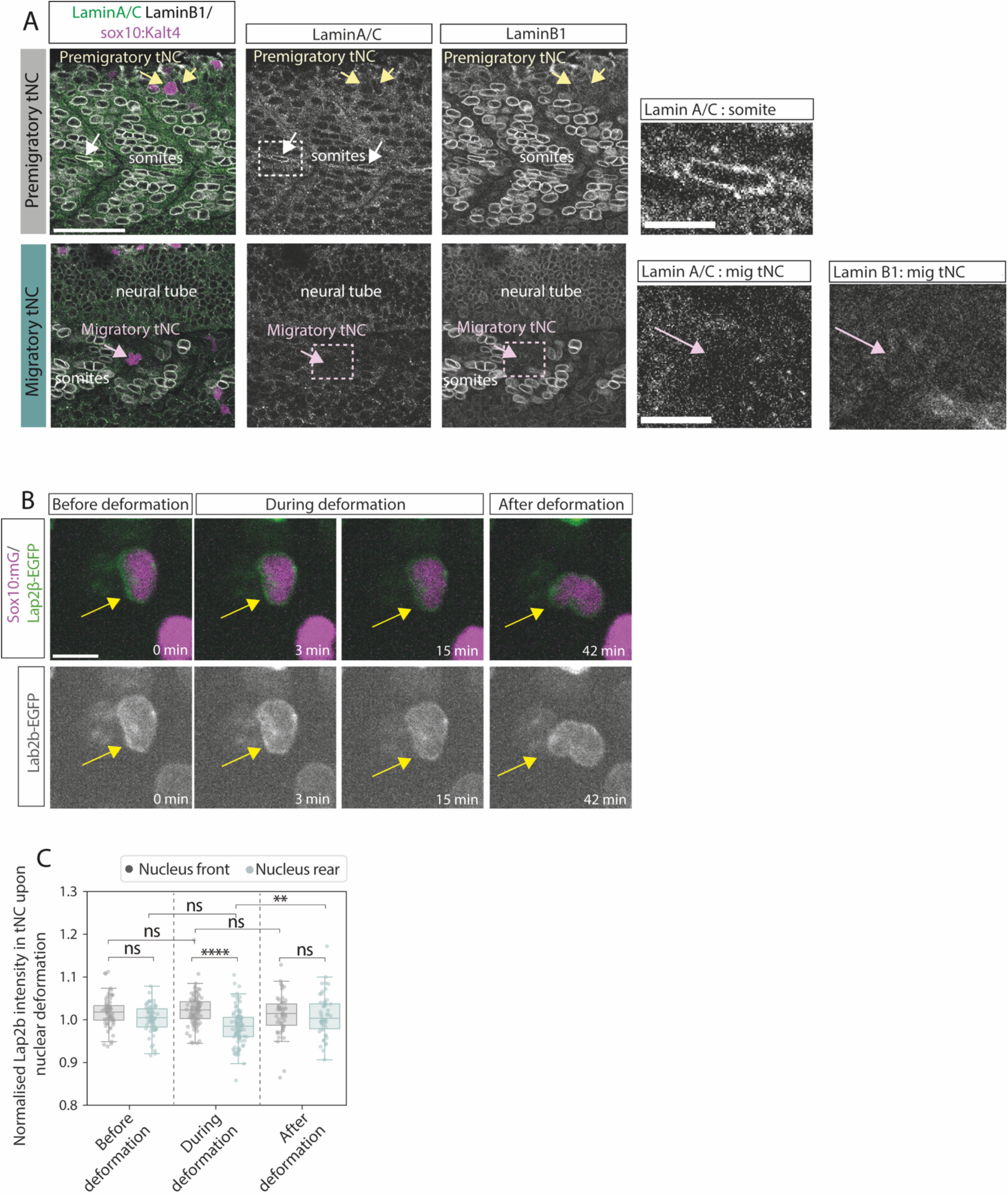
A) Representative images of expression of nuclear laminA/C and laminB1 in premigratory (yellow arrows) and migratory (pink arrows) neural crest cells and in surrounding somite (white arrows) and neural tube tissues in mid trunk region in fixed sample. Images of single Z slices. Insets (top): example of a nucleus of somite tissue expressing lamin A/C. (bottom) example of undetectable laminA/C and very low LaminB1 expression in migrating tNC nucleus. B) Representative images of Lap2β localization in the nucleus of a migratory cell in cranial region during nuclear deformation event. Yellow arrows shows the Lap2β localization at the leading front of the nucleus. C) Normalised Lap2β intensity at the leading front and the rear of migratory cranial neural crest cell nucleus before, during and after nuclear deformation event. Before deformation=n=62 timepoints from 19 cells, during deformation n=92 timepoints from 19 cells, after deformation n=50 timepoints from 19 cells. n=5 embryos. Boxplot shows mean, 1st and 3rd quartile. Each dot represents a timepoint. Scale bars 50 μm (A), 10 μm (A: insets, B).

**Movie1. Cranial neural crest migration**

Cranial neural crest migration in the region behind the eye at 10ss onwards showed in maximum projection. Overlay of membranes (magenta) and nucleus (grey) labelled tissue specifically in neural crest population using Sox10mG fish line (left) and nucleus channel alone (right) highlighting the nuclear shape changes during cell migration.

Anterior left, dorsal up.

**Movie2. Anterior trunk neural crest migration**

Anterior tNC cells transitioning from the premigratory region to migratory region under the first two somites from 12ss onwards showed in maximum projection. Overlay of membranes (magenta) and nucleus (grey) labelled tissue specifically in neural crest population using Sox10mG fish line (left) and nucleus channel alone (right) highlighting the nuclear shape changes during cell migration. Dextran labels the extracellular space (cyan).

Anterior left, dorsal up.

**Movie3. Mid trunk neural crest migration**

Mid tNC cells transitioning from the premigratory region to migratory region under two somites in the mid trunk region from 18ss onwards showed in maximum projection. Overlay of membranes (magenta) and nucleus (grey) labelled tissue specifically in neural crest population using Sox10mG fish line (left) and nucleus channel alone (right) highlighting the nuclear shape changes during cell migration. Dextran labels the extracellular space (cyan).

Anterior left, dorsal up.

**Movie4. Posterior trunk neural crest migration**

Posterior tNC cells transitioning from the premigratory region to migratory region under somite in the posterior trunk region from 30ss onwards showed in maximum projection. Overlay of membranes (magenta) and nucleus (grey) labelled tissue specifically in neural crest population using Sox10mG fish line (left) and nucleus channel alone (right) highlighting the nuclear shape changes during cell migration. Dextran labels the extracellular space (cyan).

Anterior left, dorsal up.

**Movie5. Mid trunk neural crest cells migration in spadetail mutant embryos**

Mid tNC cells transitioning from the premigratory region to migratory region in WT embryo (top) and in spt embryo (bottom). Overlay of membranes (magenta) and nucleus (grey) labelled tissue specifically in neural crest population using Sox10mG;*spt* fish line (left) and nucleus channel alone (right) highlighting the nuclear shape changes during cell migration. Dextran labels the extracellular space (grey).

Anterior left, dorsal up.

**Movie6. Neural crest migration in laser ablated embryo**

Mid tNC cells transitioning from the premigratory region to migratory region in un-ablated adjacent region (left) and in laser ablated region (right). Nuclei are labelled tissue specifically in neural crest population using Sox10mG fish line showing the nucleus shape before and after entering to migratory region.

Anterior left, dorsal up.

**Movie7. nlsGFP dynamics in migratory neural crest in trunk**

Mid tNC cell transitioning from the premigratory region to migratory region showing neural crest specific nlsGFP signal in Sox10::Kalta4;UAS-nls-emGFP fish line. Movie shows nlsGFP localisation in in cytoplasm and nucleus in before, during and after a nuclear deformation event. Anterior left, dorsal up.

**Movie8. nlsGFP dynamics in premigratory neural crest in trunk**

Mid tNC cells in the premigratory region showing neural crest specific nlsGFP signal in Sox10::Kalta4;UAS-nls-emGFP fish line. Movie shows nlsGFP localisation in in cytoplasm and nucleus. Anterior left, dorsal up.

**Movie9. BAF-cherry localisation during mid tNC migration**

Injected mApple-53BP1 trunk mRNA localisation in mid tNC in migratory region upon a during and after a nuclear deformation event. Overlay identifies the neural crest cells in green by showing the neural crest specific nlsGFP signal in Sox10::Kalta4;UAS-nls-emGFP fish line. Mid panel shows the location of neural crest nucleus. Right panel shows the homogenous BAF expression during and after the deformation event.

Anterior left, dorsal up.

**Movie10. 53BP1-dynamics in tNC during premigratory to migratory transitioning**

Injected mApple-53BP1 trunk mRNA localisation in mid tNC during premigratory to migratory transition. Overlay identifies the neural crest cells in green by showing the neural crest specific nlsGFP signal in Sox10::Kalta4;UAS-nls-emGFP fish line before cells enter to the migratory region (left panel, orange arrowheads). White arrowheads (right panel) shows the transient accumulation of 53BP1 foci.

Anterior left, dorsal up.

**Movie11. 53BP1-dynamics in tNC in premigratory phase**

Injected mApple-53BP1 trunk mRNA localisation in mid tNC in premigratory region. Overlay identifies the neural crest cells in green by showing the neural crest specific nlsGFP signal in Sox10::Kalta4;UAS-nls-emGFP fish line before cells (left panel, orange arrowheads). White arrowheads (right panel) shows the transient accumulation of 53BP1 foci.

Anterior left, dorsal up.

**Movie12. Lap2b dynamics in migratory mid tNC cells**

Injected lap2b mRNA localisation in mid tNC in premigratory region before, during and after a deformation event. Overlay identifies the neural crest cells in green by showing the neural crest specific nucleus signal in sox10-Kalt4 embryos, where chromatin is labelled with H2B-mCherry. White arrowhead shows the lap2b around the nucleus in non-deformed migratory tNC cells, white arrowheads shows the depletion of lap2b during nuclear deformation events.

Anterior left, dorsal up.

## Materials and Methods

### Zebrafish maintenance

Zebrafish were maintained and raised at 28°C on a 10h/14h dark/light cycle according to standard conditions ^1, 2^. Work was carried out under the Animal (Scientific Procedures) Act 1986 legislation under the HO Project Licence number PP8606293 (PPL holder: Dr. Elena Scarpa). Embryos were kept in 1X E3 medium (60X solution 17.2 g of NaCl, 0.76 g of KCl, 2.9 g of CaCl2·2H2O, 4.9 g of MgSO4·7). The temperature was kept at 28°C until shield stage and then adjusted between 22°C and 28°C depending on the developmental stage that embryos were to be imaged at. Prior to the experiment embryos were staged based on the morphological criteria, such as number of somites and elongation of the tail.

### Molecular Cloning

mApple-53BP1 and BAF-mCherry were subcloned into a pCS2+ vector for mRNA synthesis from Addgene plasmids 69531 ^3^and 172447 ^4^. For mApple 53BP1, pCS2+ was digested with BamHI and EcoRI (New England Biolabs) and mApple 53BP1 amplified with the following primers: 5’-CATCATGGATCCatggtgagcaagggcgag-3’; 5’-GAGGAGGAATTCctacccggtagaattatc-3’. Fragments were ligated using a T4 DNA Ligase (Promega) and transformed into competent cells. For BAF-mCherry, pCS2+ was linearised with EcoRI (New England Biolabs). BAF-mCherry was amplified by PCR using the following primer: 5’-AGGATCCCATCGATTCGAATTCatggtgagcaagggcgaggag-3’; 5’- GCTCGAGAGGCCTTGAATTCctacaagaaggcatcacaccactctcgaa-3’ and subcloned in pCS2 using the NEB Builder HiFi Kit (New England Biolabs). The 14xUAS-BGI-nucEmGFP- POUT plasmid was a gift from Harold Burgess (REF). The cmlc2-EGFP expression cassette was amplified from pDestTol2CG2^5, 6^ using the following primers: 5’- CATCATGGTACCAAAGCTTAAATCAGTTGT-3’; 5’-ATGATGGGTACCTTACTTGTACAGCTCGTC-3’ and inserted into the Kpn I site of 14xUAS-BGI-nucEmGFP-POUT using T4 DNA Ligase (Promega).

### Zebrafish transgenic and mutant lines

The following zebrafish lines were used in this study; Wild type (AB and TLF), Sox10:mG (^7^, Sox10-Kalt4 ^8^ Sox10::Kalta4;UAS-nls-emGFP ^9^, *spt-1 (b104r1)* ^10^. To generate the UAS-nls-emGFP transgenic line, 20pg of pTol1-UAS-nls-emGFP cmlc2EGFP plasmid DNA was injected with 80pg of Tol1 transposase mRNA at the 1-cell stage. F1 founders were identified by screening for green heart fluorescence. *spt-1 (b104r1)* heterozygous fish ^10^ were crossed into sox10:mG background, and Sox10mG. The primer pair, sptF (5′-TTG ACC ACA ATC CCT TTG CCA A-3’) and sptb104R (5′-GCC TTC ACC TCC AGC TCT TTA CG-3’), flanking he deletion found in sptb104 amplified approximately 0.9 kb and 1.9 kb products in the mutant and WT, respectively, and was used for genotyping ^11^. Double heterozygous fish were incrossed to generate *spt* mutant and control siblings.

### mRNA synthesis and microinjections

mRNAs were synthesized from linearised pCS2+ plasmids using the SP6 mMessege mMachine kit (Invitrogen). mRNAs were injected as follows.

#### Global labelling

Membrane GFP, membrane Cherry ^12^ and Neurocan-GFP (ss-N-Can-GFP)^13^ were injected at 1-cell stage directly to the cell with concentration of 20-50 pg/embryo, 50pg/embryo and 25pg/embryo, respectively, in WT or Sox10mG background. Embryos with the most homogenous signal were selected for experiments.

#### Mosaic labelling

mApple-53BP1 trunc and BAF-mCherry was injected into a single cell at 8- to 16-cell stage for mosaic expression into Sox10::Kalta4;UAS-nls-emGFP background. To visualize 53BP1 and BAF dynamics in neural crest cell population specifically, embryos with double expression of Sox10::Kalta4;UAS-nls-emGFP and injected mRNA were selected for mounting. Final selection of embryos with cellular co-localization of injected mRNA and Sox10::Kalta4;UAS-nls-emGFP were done at the microscope. For measurement of nucleo-cytoplasmic leakage in collectively migrating cNCs, Sox10::Kalta4;UAS-nls-emGFP were injected with 1 nl of Sox10-lynTomato plasmid DNA^14^ was injected at 1 cell stage at a concentration of 6 pg/embryo together with 30 p/embryo g Tol2 transposase mRNA.

### Visualization of the space

To visualize the extracellular space between adjacent tissues in live embryos, Neurocan- GFP mRNA injection or 10kDa Dextran red or far red was used (Alexa 546 10000 MW Dextran, Invitrogen D22911, Alexa 647 10000 MW Dextran, Invitrogen D22914). Dextran red was used for space quantification at concentration of 0.2 ng/embryo and far red was used for live imaging at concentration of 0.3-0.6 ng/embryo. To visualize the extracellular space in the cranial region NCan-GFP mRNA was injected at 1-cell stage (single time point imaging for space quantification), or Dextran-647 was injected at 1K blastula stage into the extracellular space (in vivo live imaging). For imaging the space in the trunk, embryos were injected shortly prior to experiment with dextran; embryos were mounted laterally on a 60 mm plastic dish in a single drops of 0.6% low melting point agarose (LMPA, Sigma A4018). To visualize the extracellular space in anterior and midtrunk, Dextran was injected to the extracellular space between yolk sac and EVL next to the head (Fig S2A). To visualize space in posterior trunk, Dextran was injected into the extracellular space at the tip of the yolk extension (FigS2A). Differences in extracellular space labelling and injections are justified by the different developmental stages; at earlier stages injections directly to the extracellular space between the yolk and EVL are difficult to achieve.

### Perturbation experiments

#### Laser ablation

To remove individual somites from the live embryo, presomitic mesoderm (psm) was ablated using a TriM Scope II Upright 2-photon Scanning Fluorescence Microscope controlled by the Imspector Pro software (LaVision Biotec) equipped with a tuneable near-infrared (NIR) laser source delivering 120 femtosecond pulses with a repetition rate of 80 MHz (Insight DeepSee, Spectra-Physics). The laser was set to 927nm, with power ranging between 1.05-1.20 W. The maximum laser power reaching the sample was set to 220 mW and an Electro-Optical Modulator (EOM) was used to allow microsecond switching between imaging and treatment laser powers. Laser light was focused by a 25x, 1.05 Numerical Aperture (NA) water immersion objective lens with a 2mm working distance (XLPLN25XWMP2, Olympus). Ablations were carried out during image acquisition (with a dwell time of 9.27 µs per pixel), with the laser power switching between treatment and imaging powers as the laser scanned across the sample. Embryos were treated at around 14 to 17 somite stage. mGFP positive embryos were selected and mounted in 0.6% low melting point agarose (LMPA in E3) laterally on a 60mm plastic dish in agarose wells (see section ‘Mounting embryos for in vivo imaging’). First PSM was detected by the morphology, and somites already segmented from the PSM were not targeted.The region of interest was selected by creating a ROI manually avoiding targeting neural tube or already segmented somites. The laser was set at 927nm and 85%-100% of 1.0-1.2W laser power was applied only to the selected ROI with 2µm z-interval for few µm at the time. Several rounds of ablation was performed and each round ROI was re-defined securing an accurate targeting to the PSM tissue only. The deepest PSM tissue adjacent to the neural tube and notochord (likely representing adaxial cells) was left un-treated to avoid tissue damage in the underlying structures (neural tube and notochord) and to provide guidance cues for migrating tNC . After the ablation embryos were removed from the agarose carefully and left to develop at 28°C for 2 hours and then remounted laterally in 0.6% agarose for in vivo live imaging.

#### Spadetail mutant

Double heterozygous sox10mG; *spt104R* fish were incrossed. For measurements of the extracellular spaces, embryos were injected at the 1-cell stage with X pg of Ncan-GFP mRNA. For live imaging, embryos were injected at 1000-cell stage with 10Kda Dextran-647. Embryos positive for both sox10:mG and dextran were selected and mounted for live imaging at 16 hpf as described below. *Spt* mutant embryos were morphologically identified by lack of trunk somites and by the spade-shaped tailbud. After imaging, control embryos were recovered and genotyped as previously described. Both heterozygous and *wt* siblings were included in the analysis as we did not detect differences in inter-tissue spaces or nuclear shape dynamics (data not shown).

### In vivo imaging

#### Mounting of embryos for in vivo imaging

Embryos were manually dechorionated using fine forceps and mounted on a 35 mm glass bottom dish (Ibidi, 81158) in 0.6% low melting point agarose/E3 with 0.002% MS-222 (Sigma A5040-100G). For imaging the extracellular space, embryos were mounted dorsal side up on 60mm plastic dish in agarose wells. Wells were created prior to the experiments in 1% agarose in E3 with custom made moulds with well diameter of 500 µm. For imaging neural crest cell dynamics, embryos were mounted laterally on a glass bottom 30mm dish (Ibidi, 81158). For imaging the posterior trunk, agarose was removed around the tail so that it could freely elongate. Imaging dish was filled with E3 with 0.002% MS-222.

### Microscopy

Extracellular space in cranial region was performed using a Nikon Eclipse E1000 equipped with a spinning disk unit (Yokogawa CSU10), a laser module with 491nm and 561nm excitation (Spectral Applied Research LMM2), and a C9100-13 EM-CCD camera (Hamamatsu). Image acquisition was carried out using the Volocity software (Perkin Elmer). Using a Nikon FLUOR 40X/0.80W DIC M. WD 2.0 water immersion objective with z-interval of 0.5 µm. Extracellular space in anterior, mid and posterior trunk region was imaged by using 2-photon fluorescence microscopy on a TrimScopeII multiphoton microscope, using a Ti:Sa tunable laser at 927 nm for Ncan- GFP and a 1040nm fixed wavelength laser for 10kDa Rhodamine-Dextran at 1.2W at 10- 17% laser power with 25x, 1.05 Numerical Aperture (NA) water immersion objective lens with a 2mm working distance (XLPLN25XWMP2, Olympus), using a z-interval of 0.5 µm. For live imaging of neural crest cell dynamics, we used a Perkin Elmer Ultraview Vox spinning disc microscope equipped with an incubation chamber pre-equilibrated at 28°C. Imaging was carried out with a 30X UPlanSApo Silicone immersion lens (Olympus). Usually 100μm stacks with z-interval of 1μm were acquired every 3 minutes for 8 hours with 15 to 22% laser power and 50 to 100ms exposure time for 488 and 561 channels, and 5-10% laser power and 20 to 50 ms exposure time for 633 channel. For mApple- 53BP1 trunc and BAF-mCherry live imaging was performed by using time resolution of 1 minute.

### Immunostaining

Immunostaining for nuclear antigens was carried out by adapting a previously published deyolking protocol ^15^ to improve antibody penetration. Briefly, embryos were dechorionated and fixed for 2 hours with 1% PFA in PBS, washed three times with PBS 0.1% Triton-X-100. The yolk was punctured with a fine needle and embryos were mechanically agitated with a glass Pasteur pipette to facilitate yolk detachment. Embryos were then postfixed with 4% PFA at room temperature, permeabilised with three PBS 0.5% Triton-X-100 washes and blocked at room temperature with blocking buffer (PBS 0.5% Triton-X-100, 4% normal goat serum, 2% DMSO). Primary antibodies were incubated for 48 hours at 4°C, washed in PBS 0.5% Triton-X-100, and secondary antibodies were incubated for 2 hours at room temperature in blocking buffer at a dilution of 1:500. Embryos were mounted on glass slides in 75% glycerol/PBS and imaged using either an Olympus FV3000 confocal microscope on a 30X Silicone immersion lens or a Leica SP8 confocal microscope using a 40X oil immersion lens. For collagen1a1a immunostaining, embryos were dechorionated and fixed with 4% PFA overnight at 4°C, washed and permeabilized with PBS 0.5% Triton-X-100 and blocked at room temperature with blocking buffer (PBS 0.5% Triton-X-100, 4% normal goat serum, 2% DMSO). Primary antibodies were incubated for overnight at 4°C in blocking buffer, washed with PBS 0.5% Triton-X-100, and secondary antibodies were incubated for 3 hours at room temperature in blocking buffer, washed with PBS 0.5% Triton-X-100 and DAPI was added in the final round of washes with concentration of 2.5μg/ml. Embryos were mounted on glass bottom dish laterally in 1% LMPA and imaged at Leica SP8 confocal microscope using a 40X oil immersion lens. The following primary antibodies were used: Rabbit anti-LaminB1 (Abcam Ab16048) 1:100, Mouse anti LaminB2 (Abcam AB8983), Mouse anti LaminA/C (Cell Signaling, 4777S) 1:100, Rabbit anti-gammaH2AX (Genetex IHC-00059) 1:200, Rat anti-RFP (Chromotek, 5F8) 1:200, Chicken anti-GFP (Abcam Ab13970) 1:200, CPCA anti-mCherry (EnCor) 1:200, Mouse F59 (DSHB) 1:50, Col1a1a (GeneTex, GTX133063) 1:100. The following secondary antibodies were used at 1:500 dilution: Alexa 488 goat anti-chicken (A11039,Invitrogen); Alexa 546 Goat AntiRat (A11081, Invitrogen); Alexa 546 Goat Anti-chicken (A11040, Invitrogen); Alexa 488 chicken anti-mouse (A21200,Invitrogen); Alexa 647 Fab goat anti-mouse (A48289,Invitrogen); Alexa 647 Fab goat anti-rabbit (A48285,Invitrogen).

### Sample processing for electron microscopy and SEM imaging

Samples were fixed in fixative (2 % glutaraldehyde/2 % formaldehyde in 0.05 M sodium cacodylate buffer pH 7.4 containing 2 mM calcium chloride) overnight at 4°C. Samples were then washed 5x with 0.05 M sodium cacodylate buffer pH 7.4, and osmicated (1% osmium tetroxide, 1.5 % potassium ferricyanide, 0.05 M sodium cacodylate buffer pH 7.4) for 3 days at 4oC. After washing 5x in DIW (deionised water), samples were treated with 0.1 % (w/v) thiocarbohydrazide/DIW for 20 minutes at room temperature in the dark. After 5x washes in DIW, samples were osmicated a second time for 1 hour at RT (2% osmium tetroxide/DIW), washed again 5x in DIW and block-stained with uranyl acetate (2 % uranyl acetate in 0.05 M maleate buffer pH 5.5) for 3 days at 4°C. Samples were washed 5x in DIW and then dehydrated in a graded series of ethanol (50%/70%/95%/100%/100% dry), and 100% dry acetonitrile, 3x in each for at least 5 min.

Samples were infiltrated with a 50/50 mixture of 100% dry acetonitrile/Quetol resin (without BDMA) overnight, followed by 3 days in 100% Quetol (without BDMA). Then, the samples were infiltrated for 5 days in 100% Quetol resin with BDMA, exchanging the resin each day. The Quetol resin mixture is: 12 g Quetol 651, 15.7 g NSA (nonenyl succinic anhydride), 5.7 g MNA (methyl nadic anhydride) and 0.5 g BDMA (benzyldimethylamine; all from TAAB). Samples were placed in embedding moulds and cured at 60oC for 2 days. Thin-sections (∼ 200nm) were cut using an ultramicrotome (Leica Ultracut E) and placed on melinex coverslips and allowed to air-dry. The coverslips were mounted on aluminium SEM stubs using conductive carbon tabs and the edges of the coverslips were painted with conductive silver paint. Then, samples were sputter coated with 30 nm carbon using a Quorum Q150 T E carbon coater.

Samples were imaged in a Verios 460 scanning electron microscope (FEI/Thermofisher) at 4 keV accelerating voltage and 0.2 nA probe current in backscatter mode using the concentric backscatter detector (CBS) in immersion mode at a working distance of 3.5- 4 mm. Stitched maps were acquired using FEI MAPS software using the default stitching profile and 5% image overlap.

### Image analysis

All the image analysis were performed by using ImageJ (NIH, https://imagej.net/fiji).

#### Extracellular space measurement

To quantify the extracellular space along the neural crest migratory path, dorsally acquired stacks were used. For cranial analysis, the space was measured in the region behind the eye. By using the ‘Line tool’ a 5px line was drawn across the extracellular space and intensity plot profile was created with ‘Plot profile’ function. The width of the distinguished high intensity peak was measured at every 5^th^ z-slize (2.5 µm interval) though the entire stack or until the signal was clear enough for the analysis. For the trunk region, similarly, a line was drawn across all of those somites that could be followed from the dorsal most region to the level of neural tube. Line was drawn over the extracellular space normal to the somite at the mid aspect of each somite and the distinguished high intensity peak width was measured at every 2.5 µm form dorsal to the ventral until the signal was clear enough for analysis.

#### Dynamic nuclear shape measurement

To obtain the parameters of neural crest nuclear shapes over time, maximum projected z stacks of nuclear channel from Sox10mG data sets were changed into 8-bit or 16-bit form and segmented by using 2D StarDist plugin in ImageJ. Versatile (fluorescent nucleus) model was used with percentile between 25 to 100. From the segmented data, mostly nucleus transitioning from premigratory regions to migratory regions were manually notated and tracked over time. Nucleus tracking was continued until cell divided or disappeared from the field of view. Daughter cells of the dividing cells were tracked as independent cell. Incorrectly segmented nucleus split into two or more units were corrected manually by using the brush tool with 5px size and merged together. Incorrectly segmented merged nucleus were separated when possible or discarded from the analysis. By comparing to the original 4 dimensional z-stacks with dextran labelling the extracellular space, time point when cells move under the somite was able to determine. Alternatively, bright field channel could be used to determine this time point in posterior movies. Stacks with selected and annotated nucleus were turned into RGB form and the following data processing was applied; ‘Maximum filter’ is first used with radius=1 to help to obtain the edges of the shape > ‘Find edges’ function to create outline of the nucleus shape > ‘Make binary’ function with method=MinError > ‘Skeletonize’ function to refine the edges to help a separation of neighbouring nucleus > ‘Analyze particles’ to obtain shape parameters, including circularity and aspect ratio.

#### Duration of deformation events

To compare the duration of deformation events at different anterio-posterio regions, mean circularity value of all migratory cells at all regions was first calculated (0.81) and this was set as a threshold; circularity values below the threshold were defined as a deformations. Consecutive timepoints under the threshold was defined as one deformation event and the duration of these events was calculated for each nucleus. Mean duration was calculated to those nucleus that had more than one deformation event.

#### Dynamic 53BP1 intensity analysis

To measure the 53BP1 intensity fluctuation together with nuclear shape over time, ROIs of nuclear edges obtained from the 2D StarDist segmentation (see ‘Dynamic nuclear shape measurement’ section) were applied in the 53BP1 channel and mean and maximum intensity of the signal from the maximum projected stack was obtained together with the nuclear shape parameters by using ‘Analyze particles’ function. To obtain the transient accumulation of 53BP1 signal, maximum intensity of 53BP1 was divided by the mean intensity of the nucleus at each time point.

To analyse the bright foci, mTrackJ plugin was used to manually track the foci over time.

#### Nucleocytoplasmic leakage of nls-GFP

Image analysis was carried out using Fiji (NIH: ref) Movies were corrected for Z and XY drift using the “Correct 3D Drift” plugin in Fiji. Z stacks that contained migrating neural crest streams were maximum intensity projected. Nucleus deformation events were manually identified and substacks of individual cells generated. Segmentation of the nucleus or cytoplasmic signal was carried out using the “Weka Segmentation” plugin. The nucleus mask was subtracted from the cytoplasmic binary image using the Fiji “Image Calculator” function. Binary nucleus and cytoplasm segmentation masks were converted into ROIs using the “Analyse Particle” function in Fiji, and ROIs were applied to the original movies to quantify nucleus or cytoplasm fluorescence intensity. The nucleus masks were used to measure nucleus shape descriptors using Fiji.

For cranial neural crest, cytoplasm segmentation was obtained by mosaically labelling neural crest cells with sox10-lynTomato DNA ^14^. Nucleus and cytoplasmic intensity values were normalised to their respective timepoint of higher nucleus circularity within 30 minutes of the maximum deformation event, defined as the timepoint of the minimum nucleus circularity, and cells aligned to each other by identifying the timepoint of maximum deformation.

### Lap2b-EGFP intensity profiles

Image analysis was carried out using Fiji (NIH: ref) Movies were corrected for Z and XY drift using the “Correct 3D Drift” plugin in Fiji. Z stacks that contained migrating neural crest streams were maximum intensity projected. Nucleus deformation events were manually identified and substacks of individual cells generated. Segmentation of the nucleus signal was carried out using the “Weka Segmentation” plugin.

The nucleus masks were used to measure nucleus shape descriptors using Fiji.

To quantify Lap2b-EGFP intensity profiles, the centre of the nucleus was identified manually in the 3D+t movie stack. For every timepoint of a deformation event, a 10 pixel wide line was traced from the posterior to the anterior of the nucleus along the direction of migration, and the “plot profile” function in Fiji was used to measure fluorescence intensity along the selection. Fluorescence intensity along each pixel of the profile was normalised by the mean fluorescence intensity of each line selection. Because nuclei change in length over time, individual datapoints were binned into 20 bins for each profile, where 0% represented the back of the nucleus and 100% the front of the nucleus relative to direction of cell migration. For each cell, the nucleus circularity was plotted to identify 4 phases of nucleus deformation (before deformation, during deformation, after deformation, recovery), and for the profiles of each phase were averaged together. For front/back polarization quantification, the rear peak of Lap2b-EGFP intensity was extracted by averaging the values between 5%-30% and the front peak was extracted by averaging the values between 75% and 90% for each intensity profile.

### γH2AX fluorescence intensity quantification

γH2AX immunostaining was carried out on sox10:mG zebrafish embryos. Briefly, Z-stacks were reordered as a time series using the Hyperstack>Reorder Hyperstack function in Image J. DAPI was used to segment nuclei of surrounding tissues and sox10- H2B was used to segment nuclei of NCs. Nuclear shapes were segmented using Stardist 2D. Segmented stacks were thresholded and a 1 pixel watershed filter was applied to separate nuclear masks that were in contact with each other.Using the magic wand tool, individual nuclear masks corresponding to the mid Z point of the NC nuclei were selected, added to ROI manager and annotated according to anatomical location (i.e Premig or Migratory tNC). Only one ROI was selected for each cell. For each NC nucleus, the nuclei of the immediate neighbours in the same Z stack were selected and added to manager to generate pool of Z-mached surrounding neighbours (Supplementary Methods Figure 1). ROIs were then applied to the confocal Z-stack and fluorescence intensity of γH2AX and nucleus shape descriptors were measured. The γH2AX mean fluorencence intensity for each NC nucleus of each image stack was normalised to the Z-matched surrounding tissues ROIs.

**Supplementary Methods Figure 1.**
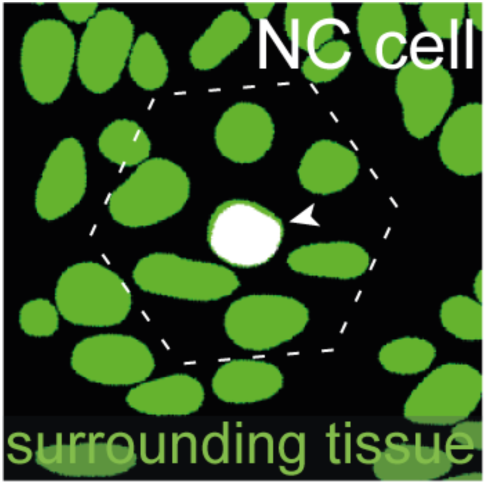
Example of NC nucleus mask (arrowhead) and its surrounding tissue neighborhood (dashed hexagonal line).

### Lamin B2 fluorescence intensity quantification

LaminB2 immunostaining was carried out on sox10:Kalt4 zebrafish embryos where H2B- mCherry is expressed under the sox10 promoter. Briefly, Z-stacks were reordered as a time series using the Hyperstack>Reorder Hyperstack function in Image J and sox10-H2B was used to segment nuclei of NCs. Nuclear shapes were segmented using Stardist 2D. Segmented stacks were thresholded and eroded 3 times. Using the magic wand tool, individual nuclear masks corresponding to the mid Z point of the NC nuclei were selected, added to ROI manager and annotated according to anatomical location (i.e Premigratory or Migratory tNC). Only one ROI was selected for each cell. Each ROI was converted in a outline of 5 pixels to detect the nuclear envelope using the ‘Make band’ function of Image J, added to ROI manager and annotated. NE ROIs were then applied to the confocal Z-stack and fluorescence intensity of Lamin B2 was measured. LaminB2 mean fluorescence for each nuclear envelope outline of each image stack was normalised to the mean grey value of the ROIs (Premigratory +Migratory tNC).

### Nuclear pore diameter measurement

Individual SEM 10kX images of trunk cross-sections (n=23 cross sections) were calibrated using the Image>Set Scale menu in ImageJ. Nuclei of Premigratory and Migratory tNC were visually identified as described in Figure S4. Nuclear pores were visually identified as gaps in the double layered nuclear envelope and nuclear pores cross-section width was manually measured using the Segmented Line tool in ImageJ.

### Statistical data analysis

The statistical tests used in each data set, p-values, the number of data points (n), and number if independent embryos for each experiment are stated in the figure legends. Statistical tests were performed using GraphPad Prism version 10.4.1. (for Windows). Data plots were generated using Python scripts (Python 3) and GraphPad Prism. In line plots mean and SEM presented. Boxplot shows mean, 1st and 3rd quartile and the whiskers extend from the box to the farthest data point lying within 1.5x the inter-quartile range (IQR) from the box.

## Notes

### Competing Interest Statement

The authors have declared no competing interest.

